# Ordered developmental emergence of number-selective neurons in days-old zebrafish

**DOI:** 10.1101/2024.08.30.610552

**Authors:** Peter Luu, Anna Nadtochiy, Mirko Zanon, Noah Moreno, Andrea Messina, Maria Elena Miletto Petrazzini, Jose Vicente Torres Perez, Kevin Keomanee-Dizon, Matthew Jones, Caroline H. Brennan, Giorgio Vallortigara, Scott E. Fraser, Thai V. Truong

## Abstract

Number sense, the ability to discriminate discrete quantity, is widespread across animals, yet how this capacity develops remains unknown. Using two-photon light-sheet imaging, we recorded whole-brain activity at single-cell resolution in larval zebrafish (*Danio rerio*) exposed to controlled visual numerosity stimuli. We discovered that number-selective neurons emerge in a striking developmental sequence: cells tuned to numerosity 1 are already present at 3 days post-fertilization (dpf), and neurons selective for 2, 3, and higher quantities appear in increasing abundance later, at 5 and 7 dpf, accompanied by a relative reduction in 1-tuned cells. The proportion of number-selective cells, summed over all tested numerosities, relative to all identified neurons in the brain, was found to decrease over time. We further showed that a machine-learning decoder based on the activity of the number-selective neurons can predict the number stimulus seen by the animals with accuracies at better than twice chance level. These results reveal how neuronal circuits develop structured numerical codes and provide a framework for studying the emergence of cognitive primitives at cellular resolution.

## Introduction

Understanding quantity, whether discrete or continuous, is fundamental for survival, such as to avoid predators, find food, mate, and conduct other group behaviors (1–8). Quantity estimation, often referred to as the Approximate Number System (ANS), allows both humans and animals to intuitively estimate numerosity, or quantity of objects in a set, without precise counting (9–11). The ANS develops during early infancy, highlighting its importance as a foundational aspect of cognition (12). It is suggested that the ANS forms the basis for complex mathematical abilities, ultimately shaping how we perceive and interact with the world (9–11). Past studies on the neural basis of number sense are limited to identifying individual neurons or specific brain regions associated with the behavior, but understanding how a network of number-selective neurons functions across the entire brain remains elusive (11,13–18). In primates, Viswanathan and Nieder (18) showed a visual sense of number mapped to the parietal and prefrontal cortices. Expanding on this, recent studies suggest that visual number processing extends beyond these regions and involves the superior colliculus, a deep subcortical area (19,20). In birds, involvement of several pallial regions has been recently documented by early gene expression (21). While the use of electrodes can access individual neurons at multiple regions, it is difficult to unbiasedly capture all neurons throughout the brain, especially without neuronal damage after implantation (22,23). Capturing neuronal activity across the whole brain would enable researchers to map neural circuits involved in numerical processing with unparalleled precision in both encoding and representation.

To address this, we developed and optimized a two-photon light sheet microscopy platform (24–27) and a customized data analysis pipeline (28) to noninvasively image the functional activity of nearly all neurons across the whole brain in larval zebrafish (*Danio rerio*) with single-neuron resolution. Larval zebrafish, which hatch and begin to utilize their visual system at 3 days-post-fertilization (dpf), offer many advantages as a model system for studying neural processes due to their transparency, genetic tractability, and drug screening applications (29–31). Using a transgenic zebrafish that expresses pan-neuronally a calcium indicator (32), Tg(elavl3:H2B::jGCaMP7f), we monitored whole-brain neuronal activity in response to non-symbolic, visual numerical stimuli. Recent studies have shown evidence of numerical discrimination in zebrafish larvae at age as early as 7 days post fertilization (dpf) (33–35). We thus reason that number-selective neurons likely are present at 7 dpf, possibly even earlier, for the animal to exhibit the number-dependent behavior. Hence, we aim to identify these number-neurons, across the whole brain in an unbiased way, to characterize the emergency of number sense during the early development window of zebrafish.

## Results

We recorded whole-brain neuronal activity of agarose-embedded zebrafish at age 3, 5, and 7-dpf, while the animals were presented with visual numerical stimuli based on dots (Fig. 1a, Methods). Identical experimental protocol, without the visual stimuli presentation, was applied to a no-stimulus control 7-dpf cohort. Imaging was acquired at a 1-Hz whole-brain volumetric rate, and the entire imaging experiment lasted for 90 minutes, as the numerical stimulus sequenced between 1, 2, 3, 4, 5 dots in pseudorandom order. When the quantity of dots changes, several non-numerical geometrical properties co-vary and can confound numerical effects. For instance, two circles have a higher combined area than a single circle of the same diameter. To minimize such confounds, the dot stimuli were designed to control for both numerical and non-numerical features (i.e., continuous physical variables that systematically vary with numerosity) (36,37). Five key geometrical covariates capture these non-numerical effects: Radius (R), Total Area (TA), Total Perimeter (TP), Convex Hull (CH), and Inter-Distance (ID) (36, 88). These covariates can be grouped into two categories: size-related covariates (R, TA, TP), which describe the dimensions of individual dots, and spread-related covariates (CH, ID), which describe the spatial arrangement of the dots. To satisfy geometric constraints, only one size covariate can be combined with one spread covariate at a time (37). Based on this rule, the covariates were combined into six feasible pairings (ID+R, ID+TA, ID+TP, CH+R, CH+TA, and CH+TP), which have also been adopted in previous studies investigating numerosity selectivity (88). The angular diameter of each dot was set to a minimum of 5°, above the zebrafish visual acuity threshold of 2–3° (38). For each stimulus presentation, a new spatial pattern was generated so that no pattern was ever repeated during the 90-minute recording session (Supplementary Fig. S1). To avoid any entrainment of neural activity to the stimulus timing, the inter-stimulus interval followed a pseudo-random sequence (Methods).

**Fig. 1.**
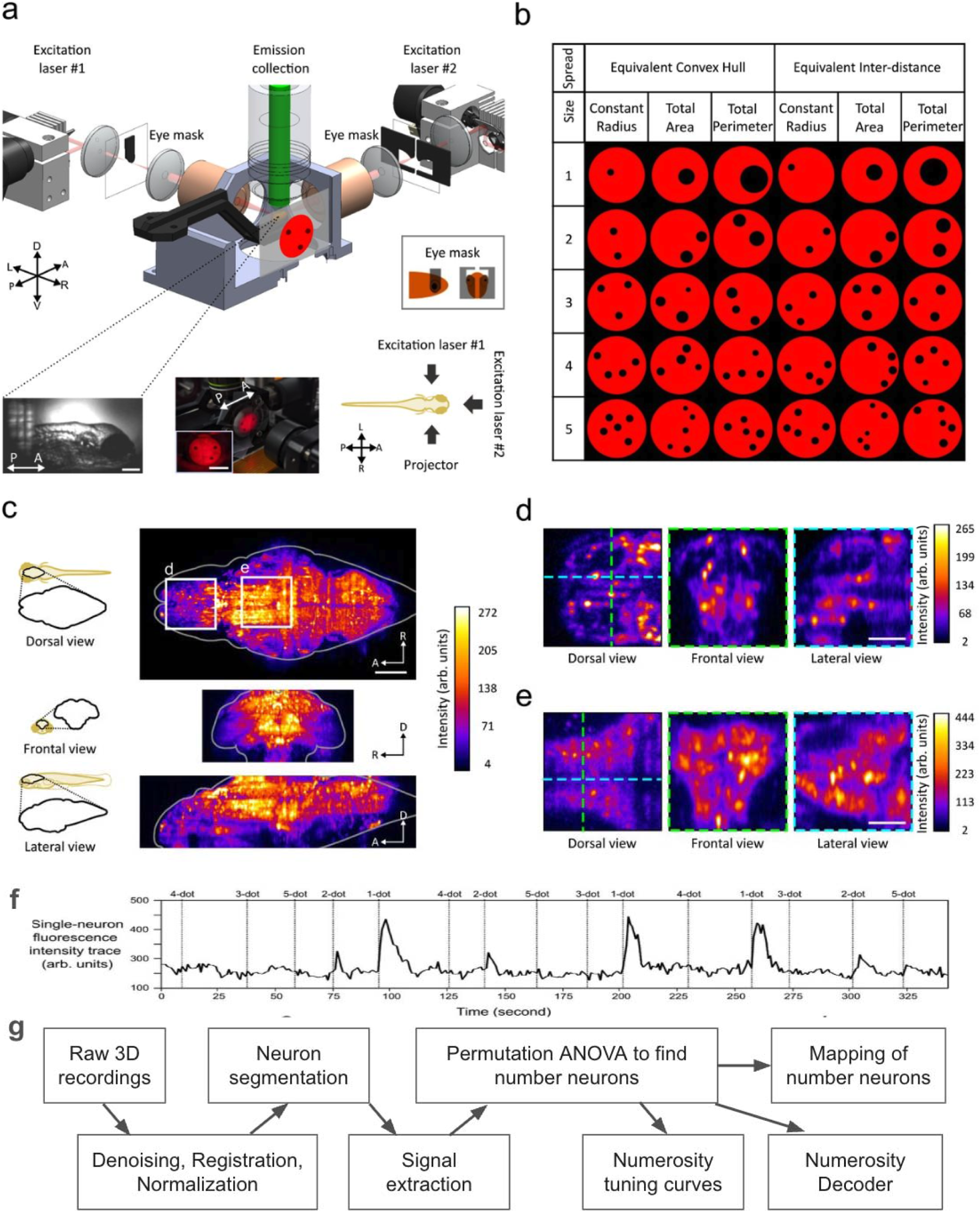
Application of two-photon fluorescence light sheet microscopy to detect neuronal representation of number perception in larval zebrafish. **a** Schematic of the two-photon light sheet fluorescence microscope. The sample is excited by two orthogonal 920 nm scanning lasers aided by physical eye-shaped masks that block laser illumination of the eyes (Supplementary materials - Materials and methods: Calcium imaging). Stimulus is displayed on the right side of the larval fish. Bottom-left: brightfield image of the mounted sample, scale bar = 200 μm. Center-bottom: stimulus projection image, scale bar = 10 mm. Bottom-right: Dorsal view of the sample relative to the stimulus direction and excitation laser. A = anterior, P = posterior, D = dorsal, V = ventral, L = left, R = right. **b** Continuous geometrical parameters used to control for non-numerical covariates when changing quantities of objects. Convex hull and inter-distance controls spread of the dots, while radius, total area and perimeter controls for the dot size. See Method: Stimuli Generation. **c** Example maximum image projection (MIP) of a 7-dpf zebrafish brain. Top, dorsal view, MIP along the dorsal/ventral axis; middle, frontal view, MIP along the rostral/caudal axis; bottom, lateral view, MIP along the left/right axis. Images were averaged over 60 seconds. Scale bar = 100 μm. **d, e** Magnified image showing cellular resolution. (d) forebrain (e) tectum from (c). Left, dorsal view of a single plane; middle, frontal view of a single plane along the green dash lines; right, lateral view of a single plane along cyan dash line. Scale bar = 50 μm. **f** Single-neuron fluorescence intensity trace of Ca^2+^ activity during numerical stimuli. Neurons responsive to numerical stimuli show a Ca^2+^ spike after the stimulus onset. **g** Schematic depicting the analysis pipeline. See texts for details.

We applied and optimized several publicly available software tools for managing, processing, and analyzing volumetric movie data and neuronal signals. We used VoDEx (28) to manage the 4D (volumetric movie) data and stimuli annotations. Advanced Normalization Tools (ANTs) (39) was used to spatially align and correct for motion artifacts in 4D datasets. Registration of multiple samples onto a representative brain template was performed using ITK-SNAP (40). To segment for the signals from individual neurons, we applied the Python toolbox for large-scale Calcium Imaging Analysis (CaImAn) (41). See Methods for full details. Fig. 1c, d, e show examples of the raw image data with whole-brain coverage at cellular resolution. Examples of the segmentation output are shown in Supplementary Fig. S2. The extracted fluorescence signal trace from a representative segmented neuron is shown Fig. 1f, showing increased amplitude in response to the numerosity stimuli. Fig. 1g depicts schematically our overall analysis workflow (Methods).

### Identification of number-selective neurons

To identify neurons specifically responsive to changes in numerosity (Fig. 2a, Fig. S3) from those responsive to geometrical changes, we applied a two-way permutation Analysis of Variance (ANOVA). Details of the ANOVA are described in Methods. Briefly, neurons were filtered based on a significant main effect for changes in numerosity (p < 0.01), without exhibiting significant main or interaction effects due to geometrical covariates (see an example in Supplementary Fig. S3). The neurons identified through the ANOVA, termed “number-selective neurons” (or “number neurons”) (13), represent the animal’s neural encoding of numerosity. We validated the robustness of the ANOVA in identifying the number-selective neurons by using a subset of the data to find these neurons, and testing their responses on the held-out data, verifying that no significant changes were found (Methods) to either the overall identity of the identified number neurons (Supplementary Fig. S4) or how they respond to the stimulus (Supplementary Fig. S5). Further, we found no time-dependent effects on the results from the 90-minute recordings (Methods, Supplementary Fig. S6).

**Fig. 2.**
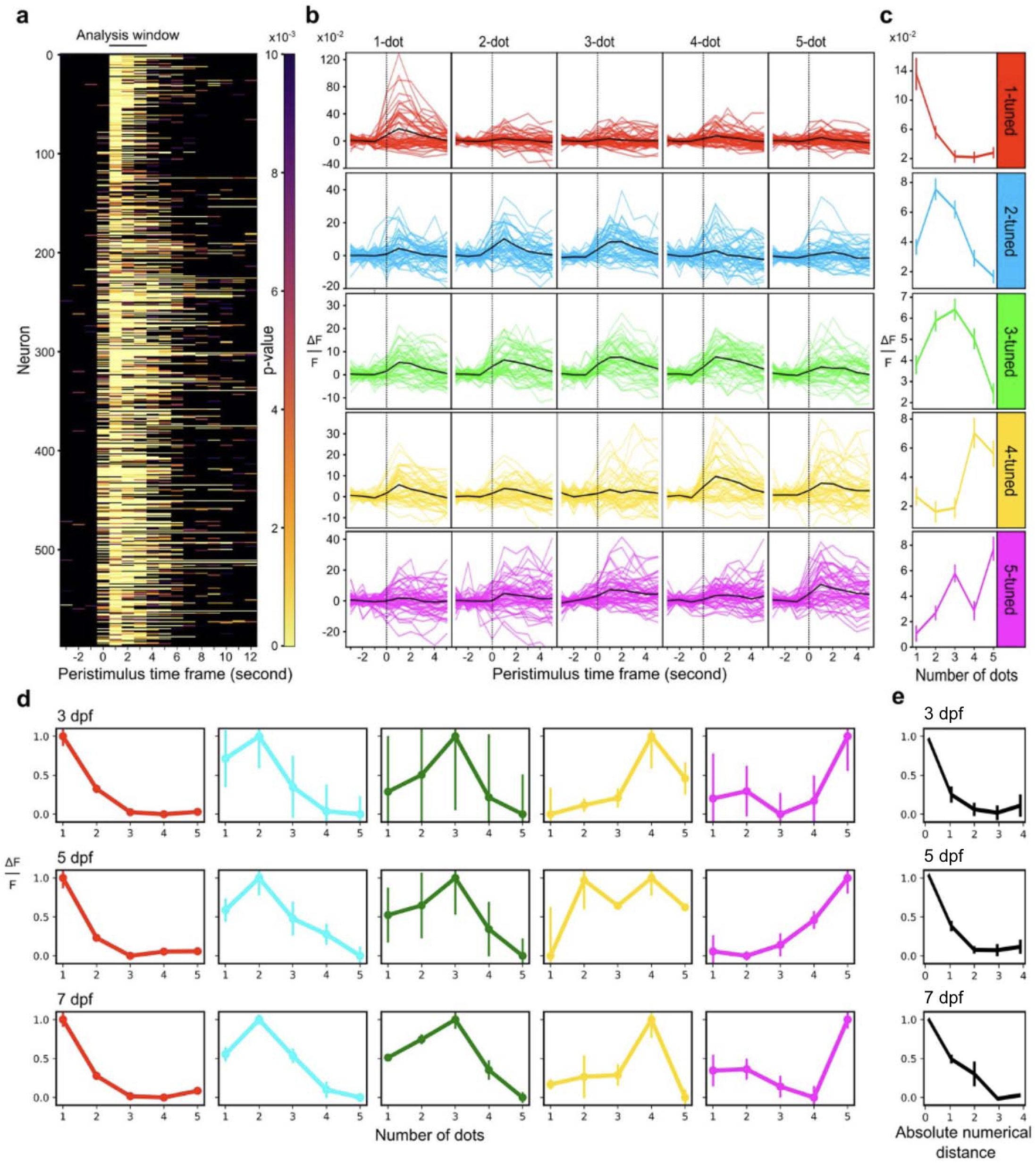
Number-selective neurons produce higher Ca2+ activity for preferred numerosities. **a** Statistical significance of number selectivity over time during stimulus onset in one example 7-dpf larvae. Rows represent individual neurons (n = 599) centered around the stimulus onset (peristimulus). Stimulus starts at time = 0 and lasts for 1 second. Black solid line on top indicates a 3-second analysis window used to detect number-selective neurons with a two-way permutation ANOVA to account for the rise and decay time of the Ca^2+^ response. **b** Ca^2+^ activity trace of 5 example neurons with preference to 1-5 dots. Traces are centered around the stimulus onset (peristimulus) for preferred and non-preferred numerosities. Neuron preferences were selected by the highest average activity for a numerosity. Top labels indicate the number of dots presented, and each color represents the specific number tuning (1 = red, 2 = cyan, 3 = green, 4 = yellow, 5 = magenta). Baseline for ΔF/F is calculated by averaging the 3 time points prior to the stimulus onset (dotted vertical line). Black line indicates the average across 48 trials. **c** Tuning curves of each of the five neurons from *b*. Each entry is the average of the 3-second analysis window for each numerosity presentation (n = 3 * 48 numerosities). Error bar = SEM. **d** Averaged tuning curves for each type of number-selective neurons (color-coded as in **c**), for each age cohort. **e** Overall differential response, averaged over all types of number-selective neurons, for each age cohort, showing the differential response between the preferred numerosity (marked by position 0) and numerosity separated from it.

The significant main effect for changes in numerosity during the stimulus onset for a representative 7-dpf larva is shown in Fig. 2a. The analysis window of 3 seconds accounts for the typical ∼2 second decay time constant of the nuclear-localized calcium signals in our samples (42). In general, a significant Ca^2+^ response to the numerical stimulus was detected 0-3 seconds from the stimulus onset. Example Ca^2+^ signal traces of neurons tuned to 1-5 objects for one fish are shown in Fig. 2b. For each number-selective neuron, the 3-second integrated response for each numerosity is used to build the numerosity tuning curve, which describes the neuron’s response across the different numerosities, peaking at the preferred numerosity (Fig. 2c) (Methods). The neuron is said to be “tuned to” or “having preference for” that numerosity.

Individual number-selective neurons’ tuning curves show a variety of behavior, from the classical profile of having a single peak at the preferred numerosity and decreasing magnitude as the presented numerosity diverged from this value, to having secondary or asymmetric peaks (see Supplementary Fig. S7 for more examples of single-neuron tuning curves). General conformation to the classical profile was observed when the tuning curves are averaged across neurons with the same preference (Fig. 2d). Averaging over all types of number-selective neurons, the overall differential response between the preferred numerosity and numerosity separated from it, as a function of absolute numerical distance, is shown in Fig. 2e (See Supplementary Fig. S8 for the analogous differential response as a function of actual (signed) numerical distance).

### Number-selective neurons develop in an ordered sequence and decrease in relative proportion over time

On average, we identified 1300±300, 800±100, 550±100 number-selective neurons, among 14000±2700, 17000±1300, 17000±2000 identified active neurons in 3-, 5-, and 7-dpf larval zebrafish, respectively (n=5 for each age group) (Supplementary Table S1-S5). Due to the varying signal-to-noise ratios across different fish and different developmental ages, subsequent quantification of number-selective neurons are based on the neuron relative proportions, which are the raw amount of number-selective neurons normalized by the total number of identified neurons, for each type of number preference, for each sample.

As the zebrafish larval age increases across 3-, 5-, and 7-dpf, we found that number-selective neurons develop in an ordered sequence: starting with a predominant proportion of 1-tuned cells at 3 dpf, neurons tuned for 2, 3, and higher numerosities appear at larger proportion later, at 5 and 7 dpf, accompanied by a relative reduction in 1-tuned cells (Fig. 3a, b). The proportion of neurons tuned to numerosity 2 or higher shows a trending increase with age (3-dpf: 4%; 5-dpf: 12%; 7-dpf: 14%) (Fig. 3a). Zooming into each numerosity-preference (Fig. 3b), we found a significant decrease of 1-tuned neurons in 3 dpf (96 ± 1%) compared to 5-dpf (88 ± 2%) and 7-dpf (86 ± 5%) groups (p < 0.05). For 3-tuned neurons, we found a significant increase in 7-dpf (5 ± 1%) compared to the 3-dpf (0.2 ± 0.1%) group (p < 0.01). Summing across all tested numerosities, we found an overall decreasing trend in the proportion of number-selective neurons, relative to the total number of identified neurons, as a function of age from 3-dpf (10 ± 2%) to 5-dpf (4.8 ± 0.7%) and 7-dpf (3.2 ± 0.6%) (Fig. 3c).

**Fig. 3.**
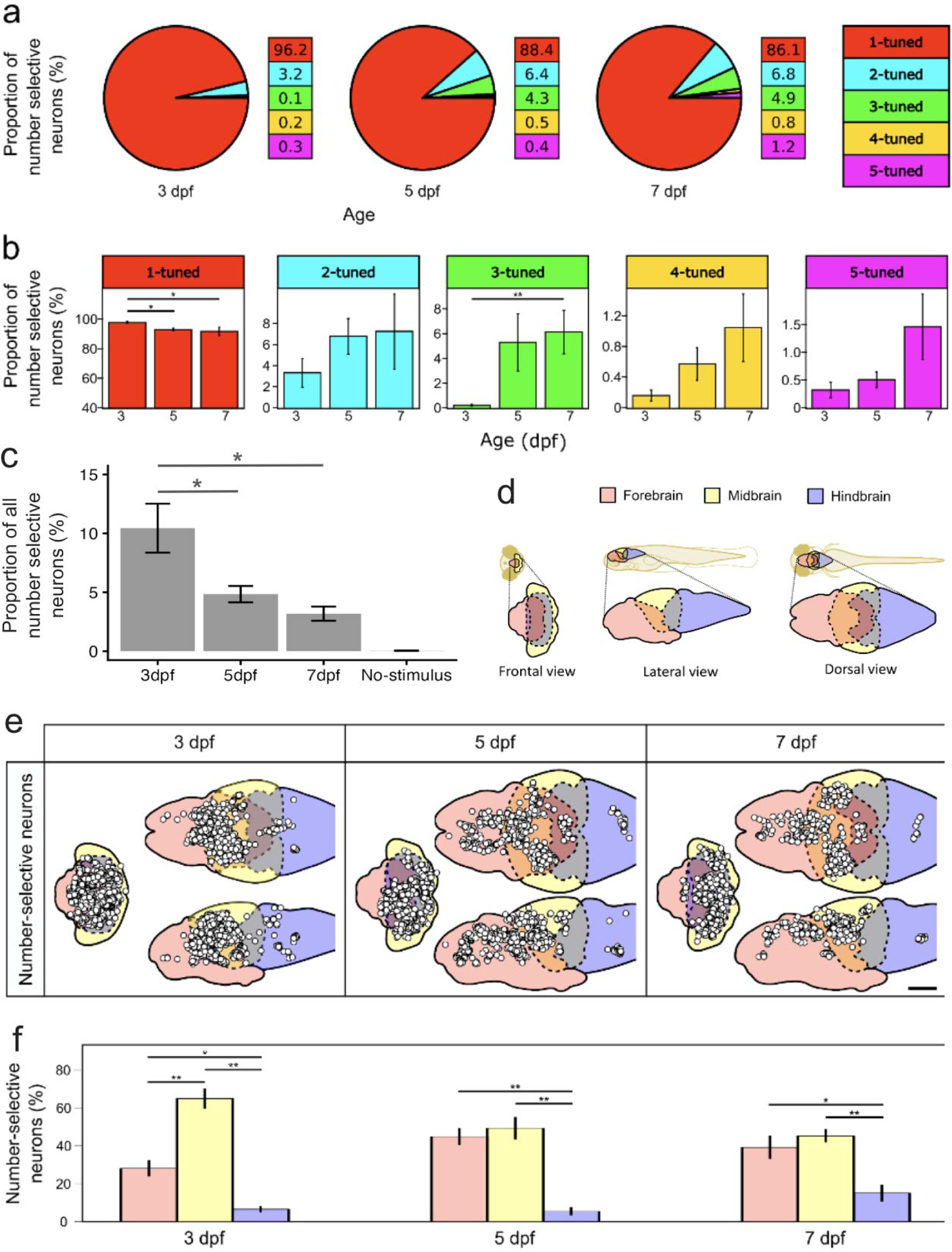
Populations of neurons tuned to specific numerosities, primarily detected in the forebrain and midbrain, show redistribution of number preference during early development. **a** Proportion of number-selective neurons as a percentage of all detected number-selective neurons across 3, 5, and 7-dpf. 1-tuned neurons = red, 2-tuned neurons = cyan, 3-tuned neurons = green, 4-tuned neurons = yellow, 5-tuned neurons = magenta. n = 5. **b** Percentages of number-selective neurons preferring specific numerosities for 3-, 5-, and 7-dpf. Percentages are expressed as a proportion to all detected number-selective neurons. Pairwise comparisons were performed using a Mann-Whitney U-test for each numerosity, multiple comparisons were adjusted using a Bonferroni correction (alpha = 0.016). Error bars = SEM, n = 5, * = p < 0.05, ** = p < 0.01. **c** Proportion of number-selective numbers, summed over all tested numerosities, relative to all identified neurons in the brain, for 3-, 5-, and 7-dpf. Pairwise comparisons were performed using post hoc parametric ANOVA. Error bars = SEM, n = 5, * = p < 0.05. **d** The 3D template of the brain was divided into three major brain regions (forebrain, midbrain, hindbrain). Solid lines indicate delineation of major brain regions; dash lines indicate overlapping regions. **e** Locations of number-selective neurons in three different individual larval zebrafish at three stages of development, representing the results as point maps in orthographic projections. The white circles represent the centers of each identified number-selective neuron. Columns indicate age. Scale bar: 100 µm. **f** Comparison of number-selective neuron distribution across brain regions of three stages of development. Number-selective neurons per region are normalized by the total number of number-selective neurons detected in the whole brain. Comparisons were performed using a two-way ANOVA for age and brain region followed by Tukey’s HSD. Error bars = SEM, n = 5, * = p < 0.05, ** = p < 0.01.

### Number-selective neurons primarily localize to the forebrain and midbrain

Number-selective neurons were detected across the forebrain, midbrain, and hindbrain of 3, 5, and 7-dpf groups. To show the locations of the number-selective neurons, we registered all imaged samples onto the respective brain templates of each age group (Methods) and mapped out the centroids of all identified neuronal nuclei (Fig. 3 c-d, Supplementary Fig. S9). We found a majority of number-selective neurons were localized to the forebrain and midbrain (Fig. 3e). For all three age groups, the forebrain (3-dpf:28 ± 4%; 5-dpf:45 ± 5%; 7-dpf:39 ± 6%) and midbrain (3-dpf: 65 ± 5%; 5-dpf: 49 ± 6%; 7-dpf: 45 ± 4%) had proportionally more number-selective neurons than the hindbrain (3-dpf: 7 ± 2%; 5-dpf: 6 ± 2%; 7-dpf: 15 ± 5%) (p ≤ 0.01). We did not identify any apparent mapping based on preferred numerosities (Supplementary Fig. S10). In the 3-dpf group, we found significantly less proportion of neurons in the forebrain than the midbrain (p < 0.001).

Building upon these findings, we further investigated the distributions of number neurons in the subregions of the forebrain (eminentia thalami, hypothalamus, pallium, pretectum, thalamus, subpallium) and found no significant changes with age (Supplementary Fig. S11, Supplementary Table S6).

### Number stimulus can be decoded from number-selective neurons

To determine if the Ca^2+^ activity of number-selective neurons across the brain is sufficient to predict the correct number of dots shown during a visual stimulus, we built a support vector machine (SVM) supervised decoder to predict the visual stimulus from the recorded activities (45). An independent decoder was built for each fish, where the features are the single-neuron Ca^2+^ activities of the entire ensemble of identified number-selective neurons (Fig. 4a, Methods). For each fish, the decoder was trained on a subset of the data and tested on the held-out data, in predicting the five presented stimuli (1, 2, 3, 4, or 5 dots). The performance of the SVM decoders, averaged for all fish for each age cohort, are described in the confusion matrices in Fig. 4b, c. (For the decoder performance for each individual fish, see Supplementary Fig. S12.) In these matrices, the horizontal axis lists the numerosity stimulus that the encoder predicts that the animal is being presented with, while the vertical axis depicts the actual experimental stimulus. Each entry of the matrix’s diagonal shows correct prediction of the true labels. The no-stimulus dataset yielded prediction accuracy of about 20% for all numerosities, consistent with chance level (1 out of the 5 tested numerosities). For all three age cohorts, the prediction accuracy was well above chance level for all numerosities (Fig. 4b): 3dpf 1 (45%), 2 (39%), 3 (37%), 4 (40%) and 5 (49%); 5-dpf, 1 (53%), 2 (44%), 3 (45%),4 (38%) and 5 (41%); 7-dpf, 1 (45%), 2 (38%), 3 (40%), 4 (36%), and 5 (45%). By averaging the diagonal entries of the confusion matrices, the decoder accuracy across all numerosities can be obtained, yielding more than twice better than chance for all age cohorts (Fig. 4c).

**Fig. 4.**
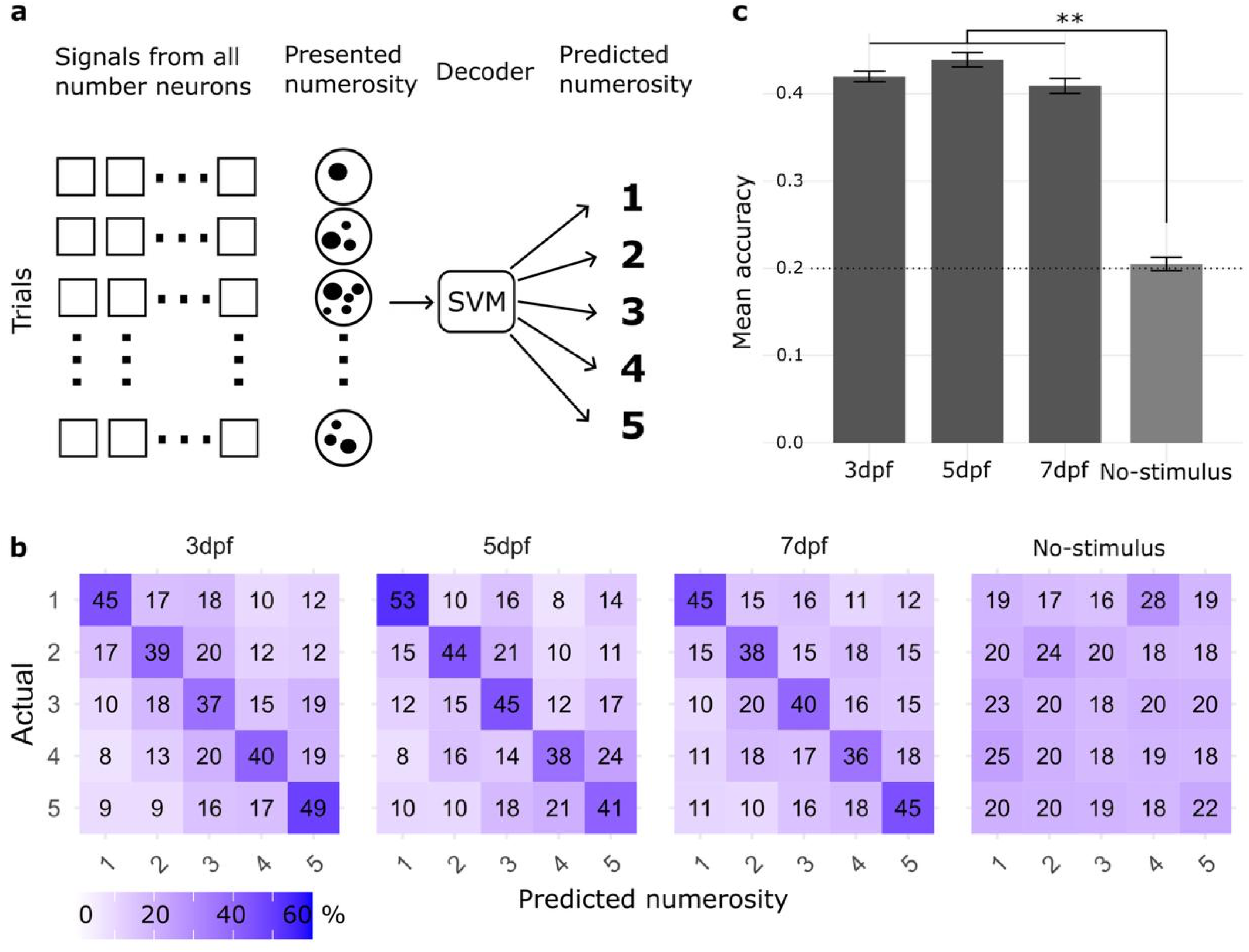
Prediction accuracy of the numerical stimuli from Ca2+ activity of number-selective neurons with support vector machine (SVM) decoder. **a** Schematic depicts the design of our SVM decoder. Feature extraction of Ca^2+^ activity from numerically-tuned neurons (average fluorescent signal in the 3-s from stimulus onset). The visual stimulus (1, 2, 3, 4, 5 dots) serves as the true label for each instance (trial). Each sample zebrafish is comprised of 240 instances (48 repetition * 5 visual stimulus). Cross validation was performed using a leave-one-out cross validation, where training was performed on the data from seven out of eight supertrials, and test was run on the remaining supertrial. Results are then average over the eight possible 7+1 splits and over the 5 fish in the age group. See Methods for details. **b** Confusion matrix of the SVM decoder of 3, 5, and 7-dpf age groups and a no-stimulus 7-dpf group. Matrix entries correspond to percentages, defined as the ratio of the number of correct predictions over the total instances of each true label. Random chance = 20%. **c** Overall SVM classification accuracies for the three age groups. Dash line indicates chance level at 20%.

## Discussion

In this study, we investigated the neural basis of number sense in larval zebrafish, focusing on the tuning of neurons to specific visual-based numerosities under different conditions. Our work has, for the first time to our knowledge, discovered the existence of number-selective neurons in larval zebrafish. The fast volumetric imaging rate of one volume per second and the high signal-to-noise ratio of light sheet microscopy enabled whole-brain recording and segmentation of approximately 17,500 active neurons per larva. The use of two-photon excitation allowed for increased depth of coverage (deeper imaging) and reduced visual artifacts compared to one-photon excitation (24,25,27,46).

To obtain robust statistical power, we adopted a recording duration of 90 minutes, balancing the need for sufficient trial numbers with the minimization of confounds such as photobleaching or habituation. While some reduction in signal amplitude may occur over time, this would only lower detection sensitivity (increasing false negatives) without introducing false positives, and animals consistently remained in good condition after recordings. Post-hoc analysis also confirmed that there were no time-dependent confounding effects found in our results due to the 90-minute recording time (Supplementary Fig. S6).

Consistent with studies using chick models (47) and human infants (12,48), we detected number selective neurons during early post-embryonic stages in zebrafish at 3 dpf (equivalent to post-natal in mammals), essentially as soon as the visual system starts to function (49). Notably, the existence of these number-selective neurons at 3 dpf precedes any known numerically-driven behaviors such as hunting and shoaling (which start at 5 and ∼24 dpf, respectively) (33–35,50), underscoring the fundamental role and necessity of early numerical cognition for survival.

Weber’s law, a foundational principle in psychophysics, states that the ability to discriminate between two stimulus magnitudes depends not on their absolute difference, but on their ratio (51). In the domain of numerical cognition, this principle is captured by the approximate number system (ANS), which encodes quantities with a precision that scales with magnitude. A hallmark of ANS representations is that tuning to numerosity is broad: neurons or populations respond most strongly to a preferred numerosity but also to nearby ones, with the width of this response increasing for larger numerosities. Consequently, tuning curves (particularly when averaged at population level) are not expected to be perfectly sharp but instead show graded, overlapping activity that reflects Weber’s law.

In line with this, the tuning curves we obtained exhibit broad profiles (see Figure 2c for an example of a single neuron - with additional examples in Supplementary Figure S7, and Figure 2d for the population average for each type of number-selective neuron), supporting the view that the underlying coding of numerosity relies on approximate, ratio-dependent representations rather than exact, discrete one, or monotonic one. This is confirmed when plotting the average response to different numerosities against the absolute numerical distance between the ‘preferred numerosity’ and the presented one, showing a decreasing response profile as the distance from the preferred numerosity increases (Fig. 2e). On the other hand, at the single-neuron level, many neurons showed asymmetric or secondary peaks (Supplementary Fig S7). Such variability has also been reported in other species (13,52) and likely reflects both developmental dynamics in zebrafish larvae and the inherent diversity of neural coding strategies within the population. Notably, the average tuning curves appear to become less noisy as the animals develop from 3 to 5 and 7 dpf (Fig. 2e). While we cannot exclude the contribution from purely experimental factors, this trend suggests that number neurons fine-tune their responses with age.

We observed a progressive, ordered emergence of number-selective neurons: at 3 dpf nearly all selective cells are tuned to 1, then cells that are tuned to 2, 3, 4, and 5 appear in increasing proportion for 5 and 7 dpf, accompanied by a reduction in the proportion of 1-tuned cells (Fig. 3a, b). This development pattern indicates a progressive, ordered construction of numerical representations, where selectivity for larger numerosities builds upon earlier-established codes. An interesting question for future studies is whether this increase is due to the generation of new neurons preferring higher numerosities or the re-tuning of existing neurons.

Together with the sequential emergence of number-selective neurons, we found that the proportion of all number neurons (summed over all tested numerosities), relatively to all identified neurons in the brain, decreases with developmental age from 3 to 7 dpf, by more than 3-fold (Fig. 3c). This result indicates a developmental pruning process where the quantity of number neurons, relatively to the total number of neurons in the brain, gets lower as the zebrafish gets older. In conjunction with the finding discussed earlier that the average tuning curves of number neurons get less noisy with age (Fig. 2e), our results suggests that as the zebrafish gets older, their number sense depends on fewer number-selective neurons, but with more refined tuning of these neurons.

The proportion of number-selective neurons we identify (∼ 5–15% of segmented cells) may appear high at first glance. However, it is important to note that our segmentation approach captures only a subset of the full neuronal population, both because the promoter elavl3 in fact does not label all neurons (∼75% coverage at best (53)) and because we restricted analyses to only neurons active during the stimulus window, thereby biasing toward task-responsive cells. Independent estimates suggest that the true neuronal population at 3–7 dpf comprises ∼ 50,000–100,000 cells (54), implying that our identified number-selective neurons represent only ∼ 1–3% of all neurons in the larval brain. This proportion is in line with, or even lower than, percentages reported in other species (e.g., ∼20% in crows (13); ∼11% in chicks (47)). Our study provides the first whole-brain estimate, unlike prior electrophysiological work confined to restricted regions, which may partly explain differences in prevalence. We interpret these results as consistent with early developmental stages in which neural representations of numerosity emerge before being consolidated into behavior.

The increased proportion of neurons preferring larger numerosities (>2) developing after 3-dpf may be caused by an improvement in visual acuity rather than changes to number-selective neurons. However, the zebrafish eye is emmetropic at 3-dpf (49), and no differences in visual acuity were detected when comparing larvae at 4-, 5-, and 6-dpf (38). Furthermore, recognition of 5 dots does not require finer visual acuity than 2 dots when maintaining equivalent inter-distances (Fig. 1b). Because we detected neurons preferring 2 dots in 3-dpf larvae, the increase of neurons preferring larger numerosities in older larvae is unlikely to be caused by improved visual acuity.

We identified number-selective neurons localized throughout the forebrain and midbrain (Fig. 3d). In the 3-dpf group, we found significantly less neurons in the forebrain compared to the midbrain, whereas in the 5-and 7-dpf groups, the forebrain contained a similar proportion of these neurons (Fig. 3e), likely due to the forebrain being more developed in the older fish (55). Further analysis of the forebrain subregions found no significant age-related changes (Supplementary Fig. S11, Supplementary Table S6). These results suggest that while overall development affects neuron distribution, the specific sub regional changes may not happen until the brain matures beyond 7-dpf.

In the mammalian brain, most number processing is to our knowledge localized to the prefrontal and parietal cortices (11,52). In non-mammals, such as zebrafish, the pallium generally fulfills the role of the prefrontal cortex (56). However, the non-mammalian brain lacks a structure that is directly analogous to the parietal lobe; instead, the optic tectum of the midbrain serves many cortical functions such as sensory processing and spatial perception (57–59). In line with our result of finding number-selective neurons in the midbrain, emerging studies suggest subcortical and optic tectal involvement in visuospatial processing on numerosities (4,19,60,61). This suggests that the optic tectum participates in more complex functions than its long-studied roles in visual mapping and sensory integration.

To assess the predictive capabilities of the number-selective neurons across 3, 5, and 7-dpf zebrafish, we trained a supervised decoder using their underlying Ca^2+^ activity. The SVM decoder achieved prediction accuracies well above chance across all developmental stages (Fig.4b-c), demonstrating that the Ca^²⁺^ activity of number-selective neurons contains reliable information about the numerosity of the presented stimulus. This consistent performance across 3-, 5-, and 7-dpf larvae indicates that a robust population code for numerosity is already established early in development and remains stable as the visual system matures. Notably, the significant decoding accuracy at 3 dpf, when the visual circuitry is only beginning to form, suggests that numerical information may be an intrinsic feature of early sensory processing rather than a product of extensive experience, pointing to the fundamental nature of number sensitivity in the developing brain.

Across the age groups of 3, 5, and 7 dpf, the SVM decoder predicted the experimental numerosity stimuli with similar accuracy (Fig. 4c). Ostensibly, this counters the intuition that the decoder should work better with older animals since the numerosity tuning curves were found to be less noisy for the older age groups (Fig. 2e). We note that on average younger zebrafish have many more number neurons, in absolute numbers, than older fish (∼ 1300, 800, and 550 neurons, for 3, 5, 7 dpf, respectively). Thus, it is possible that the higher abundance of number neurons, in the younger animals, could compensate for their noisier tuning curves to produce the statistical significance for the decoder to arrive at similar prediction accuracies across the age groups. Further, as have been generally recognized (see e.g. (62)), the performance of decoders based on an animal’s experimentally measured neural signals should not be equated to the animal’s subjective experience, as there is no guarantee that the animal would utilize the same set of neural signals or follow the identical computational model of the decoder. Thus, the reported prediction accuracies should not be interpreted as representing the actual numerical discrimination ability of the zebrafish. Rather, the success of the decoder in predicting the numerosity stimuli serve to confirm that the ensemble activity of the identified number neurons possesses robust numerical information.

One limitation of our study is the multi-prong challenges that the developing zebrafish brain presents to the imaging-based identification and comparison of neurons, from 3 to 7 dpf. First, as the zebrafish brain matures, it becomes bigger and more challenging to image optically, leading to an overall reduction in the whole-brain image quality, which ultimately results in *less* neurons being successfully identified for older animals (all else being equal). Second, the expression level of the elavl3 protein, which is used as the promoter to drive the expression of the calcium indicator, does decrease over time (53), resulting in *less* neurons being identified for older animals. Third, the zebrafish brain gains more neurons as it develops from 3 to 7 dpf (approximately going form 50,000 to 100,000 neurons (54)), resulting in *more* neurons being identified for older animals. To mitigate the effects of these challenges, we have used, as the main quantitative figure in our analysis, the relative proportion of number-selective neurons per each experimental animal, which is less sensitive to variability than absolute cell counts. Nevertheless, we recognize that the relative proportion metric does not fully negate the possible confounding factors discussed above. Thus our conclusions about the development of number neurons should be further confirmed in future studies that enlist additional technical improvements to overcome the challenges of imaging a developing brain, such as using sophisticated strategies to ensure long lasting pan-neuronal gene expression (63) and brighter calcium indicators (64).

One particularly relevant future direction is to employ the longitudinal imaging approach, in which the same larva is repeatedly imaged, released, and re-imaged across developmental stages. This would provide a robust means to identify and track the identity of the number cells over time, providing more direct insights into developmental dynamics and coding stability. To realize this longitudinal imaging approach, our experimental platform would have to be refined in several key areas, such as maintaining zebrafish viability through repeated mounting and dismounting rounds, and the development of a framework to follow neurons in a growing brain. We view this as an important direction for future research.

Overall, our findings contribute to the growing understanding of the developmental stages of cognitive abilities in vertebrates, offering new insights into how early neural circuits involved in numerosity evolve even before behaviorally measurable traits emerge. The identification of number-selective neurons in larval zebrafish as early as 3-dpf, well before the onset of numerically-driven behaviors, emphasizes the critical role of early neural development in the establishment of cognitive functions necessary for survival. This study not only adds to the foundational knowledge of numerical cognition in non-mammalian species but also opens avenues for comparative studies across vertebrate models, including primates and humans, to explore the evolutionary conservation of these neural circuits.

## Methods

### Animal care

Casper zebrafish (*Danio rerio*) expressing a pan-neural, nuclear-localized fluorescence Ca^2+^ reporter (elavl3:H2B::jGCaMP7f) was a gift from the lab of David Prober (California Institute of Technology). Following established protocols (65), zebrafish larvae were raised in 50 mL petri dishes with approximately 50 larvae per dish, in E3 medium (5 mM NaCl, 0.17 mM KCl, 0.33 mM CaCl2, 0.33 mM MgSO4), subjected to 13-hr:11-hr light:dark cycle starting at 1 day post fertilization (dpf), and fed dry food twice daily starting at 5 dpf. Experiments used zebrafish ranging from 3 to 7 dpf. Sex is not defined at this stage of development. All animal procedures conformed to the institutional guidelines set by the University of Southern California Department of Animal Research.

### Calcium imaging

Zebrafish larvae were embedded in 2% low-melting-point agarose (Invitrogen Cat 16520100) and mounted in a custom sample holder, then acclimated in the temperature-controlled sample chamber, at 28°C, for at least 30 minutes before imaging. The sample chamber was continuously perfused with oxygenated water using a peristaltic pump. Image acquisition was performed on a custom-built two-photon light sheet microscope (26) that was further modified to increase the nonlinear fluorescence signal, by optimizing the excitation polarization and implementing laser switching between the front and side light sheet excitation arms (27,66). Two-photon excitation was carried out with a Chameleon Ultra II Ti:Sapphire laser (Coherent) at 920 nm, with approximately 180 mW average power at the sample (summed from both the front and side light sheets). Excitation light sheets were delivered through a pair of air objectives (5x, 0.10 NA, Olympus LMPLN5XIR LWD M PLAN). Emitted fluorescence was collected with a water-dipping objective (20x, 1.0 NA, Olympus XLUMPLFLN-W), bandpass filtered (525/45 nm, Semrock FF01-525/45-25-STR) and recorded with a scientific complementary metal-oxide semiconductor (sCMOS) camera (Hamamatsu ORCA-Flash4.0 V3). Continuous images were acquired at a rate of 1 volume per second, in which a volume is composed of 60 z-slices at 540 x 296 pixels, over a volume of ∼ 500 x 900 x 230 (xyz) µm^3^, yielding a voxel size of 1.68 x 1.68 x 3.83 (xyz) µm^3^. Software control and hardware synchronization for image acquisition was performed as previously described (26) using µManager (67) and LabVIEW (National Instrument).

### Stimuli Generation

Dot patterns were generated using GeNEsIS (37) and controlled for convex hull, inter-distance, total area, total perimeter, and radius in (Fig. 1b). Convex hull describes the smallest convex polygon that encloses all of the visual elements. Inter-distance is the average distance between the elements. Total area equates average brightness and cumulative surface area for different numerosities. Total perimeter equates the cumulative circumference of all the elements for different numerosities. The parameters are summarized in Supplementary Table S8. Note: parameter values are applied by use case (“1” dot stimuli do not have convex hull or inter-distance parameters, constant radius does not use radius variability). Angular diameters of dots were kept above 5° to maintain visibility (38) and below 18° to avoid an escape response (68). Numerical dot elements were colored black on a (luminous) red background to avoid producing light with wavelength in the blue-green range that would interfere with the fluorescence measurements. We chose black dots over red background, rather than red dots over black background, to minimize the change in luminosity between the numerical patterns and the inter-stimulus blank red background (as the total area of the dots is smaller than that of the background for any of the used patterns).

### Visual number-based display

Visual stimuli were projected onto a diffuser placed 19 mm away from the larvae, facing its right eye from the animal’s right side (Fig. 1a). The diffuser is made of cellulose acetate (Scotch Magic Tape), with its surface normal orienting orthogonal to the zebrafish’s sagittal plane. Illumination was generated using a Qumi Q5 LED Projector (Vivitek) and bandpass filtered at 660/45 nm (Thorlabs). The stimulus was displayed for 1 second followed by varying inter-stimulus intervals of a blank red background between 15-30 seconds. The display area was 22 mm in diameter or 66° in angular diameter. Stimulus control was performed using the PsychoPy toolkit (69).

Each experimental recording has 240 stimuli presentations, or trials. These presentations are grouped into 8 supertrials, which represent a complete round of presentations consisting of all 6 geometrical control combinations, for 5 numerosities, making up 6 x 5 = 30 stimuli presentations in each super-trial. These 8 super-trials are independent repeats, and a distinct stimuli pattern is used in each of the presentations. For the entire recording, each numerical stimulus is presented 48 times following a pseudo-random order, meaning the alternation of the control conditions was random but repeated in a loop (Supplementary Fig. S1).

### Cell segmentation

The image datasets were first corrected for camera noise by subtracting a background image recorded with the fish sample, then motion corrected using Advanced Normalization Tools (ANTs) (39). Cell segmentation was performed using the Python toolbox Calcium Imaging data Analysis (CaImAn) (41). CaImAn, an established and widely used tool in the field (70,71), uses constrained non-negative matrix factorization to decompose the Ca^2+^ image data into spatial and temporal components to segment individual neurons and extract their activity. Prior to cell segmentation, the data size was reduced by selecting only the time points around the stimulus presentation (3 s before stimulus + 1 s stimulus + 5 s post-stimulus; see Fig. 2). The final volumetric time series was reduced from 5,472 s to 2160 s (9 s window x 5 numerosities x 48 repetitions). Large data handling and annotation were managed using the VoDEx Python library that facilitated time annotation and image processing (28).

The segmentation was performed in 2D where each time point consisted of 60 z-slices with the following parameters: ‘decay_time’ = 5 (length of a typical transient in seconds); ‘gSig’ = 3×3 (expected half size of neurons in pixels); min_SNR = 1.5 (signal to noise ratio to accept a component); rval_thr = 0.85 (space correlation threshold to accept a component). The ‘K’ parameter is the expected number of cells to be segmented that serves as a starting point for optimization. Since the number of cells expected in each z-slice varies, the ‘K’ parameter was estimated based on the standard deviation of Ca^2+^ fluorescence signal in each z-slice over time. The standard deviation image was thresholded at (min_std + 0.08*(max_std - min_std)) resulting in an image with pixels that represent cells with Ca^2+^ signal changes. The maximum number of the resulting pixels were then divided by ‘gSig’ to approximate the number of cells per z-slice. Since a single cell’s Ca^2+^ signal can span across 2-3 z-slices, we eliminated duplicates by merging cells with both a centroid distance less than 1 pixel and with a Ca^2+^ activity correlation coefficient higher than 0.95.

To remove false positive segmented cells (e.g. cells that were found outside of the brain), we calculated the ratio of the standard deviation to the mean, for each timepoint in the peristimulus windows. We found that a ratio of 0.05 was sufficient to remove false positive cells outside of the brain. Baseline fluorescence (F_0_) was defined as the average over three time points (i.e. 3 s) before the visual stimulus presentation for each numerosity, and the fluorescent signal (F) for each trial was normalized accordingly as (F-F_0_)/ F_0_.

### Number neuron identification with ANOVA

To identify number-selective neurons, we applied a two-way permutation ANOVA in the form Response ∼ Numerosity * Control (see (72) for a full description of this statistical analysis approach). Briefly, our data consisted of the factors Numerosity (5 levels: 1 to 5 dots) and Control condition (6 levels: ID+R, ID+TA, ID+TP, CH+R, CH+TA, CH+TP; where: CH=convex hull, ID=inter distance, R=radius fixed, TA= total area, TP=total perimeter). These six combinations of the five covariates represent the only geometrically feasible pairings that allow simultaneous control of more than one non-numerical covariate while still permitting systematic variation in numerosity, ensuring maximal independence between numerical and geometrical parameters. Given that each combination was repeated 8 times, we have 48 data points per numerosity for each neuron. For the permutation procedure, we first computed the real F values for both factors on the original, unshuffled data. Then, to generate a null distribution, we randomly shuffled the trial labels (keeping the neural activity unchanged, preserving the structure of the dataset) 10,000 times and recalculated the F statistics for each shuffle. This produced a distribution of F values as expected by chance, and the permutation p-value was defined as the proportion of shuffled F values that exceeded the real F value. Neurons were classified as number-selective if their real F for numerosity exceeded the 99% of values under the null distribution (alpha = 0.01), while all other contributions (control factor and interaction) were non-significant ensuring that detected selectivity was not driven by random fluctuations in trial identity nor control conditions. See Supplementary Fig. S3 for a representative example of a neuron that significantly responded to numerical information but not geometrical covariates.

To further validate the identification of the number neurons and assess the consistency of results across independent subsets of data, we implemented a leave-one-out (LOO) cross-validation framework for the permutation ANOVA. For each zebrafish sample, the data were divided into two groups: an *Analysis* group comprising 7 of the 8 supertrials (each consisting of 5 numerosity × 6 control conditions), and a *Testing* group containing the remaining LOO supertrial. The two-way permutation ANOVA was performed on the Analysis group to identify number-selective neurons. This procedure was repeated eight times, each time leaving out a different supertrial, thereby yielding eight independent sets of identified number neurons. These were then used in two tests to evaluate the reliability and robustness of the number neurons identification procedure. First, we quantified the overlap in the identities of the number neurons obtained from each pair of Analysis groups. For each numerosity, we computed the proportion of neurons that were common between two groups relative to the total identified neurons, yielding an overlap percentage. The overlap percentages were consistently higher than the chance level of 50% for all experimental groups (p < 1.05 × 10 ³², one-sided one-sample t-test for the average overlap ratio in each age cohort), while remaining at chance level for the no-stimulus control cohort (p = 1.00, one-sided one-sample t-test) (Supplementary Fig. S4). In the second test, we used the LOO Testing group to assess the functional reliability of the number neurons identified from the Analysis group. Specifically, we compared the tuning curves, five-component vectors describing neural responses across the five numerosities (1–5; see Fig. 2c i), computed from the Analysis (t_A) and Testing (t_T) groups for each identified neuron. We then calculated the Pearson correlation coefficient between t_A and t_T, termed the Tuning Correlation, to provide a quantitative measure of how similar the tuning curves are to each other, and thus demonstrating the tuning stability across independent data subsets. Corresponding analyses were also performed on shuffled datasets, in which numerosity labels were randomized, to obtain chance-level Tuning Correlation distributions. We found that the averaged Tunning Correlation between the experimental Analysis and Testing groups were significantly higher than the scores from shuffled data, while the no-stimulus data (which should contain no number neurons) yielded no significant difference with shuffled data (Supplementary Fig. S5). This demonstrates the numerosity tuning curve stability of the identified neurons, when tested on independent data (which were not used in identifying them). Taken together, these additional LOO cross-validation testing provide quantitative support for the reliability and robustness of our analysis pipeline in identifying the number neurons.

### Constructing numerosity tuning curves

Tuning curves were computed to characterize the numerosity selectivity of individual neurons and to derive population-level representations. At the single-neuron level, the tuning curve was defined as the mean normalized ΔF/F response of the neuron during the 3-s period following stimulus onset, calculated separately for each of the five numerosity conditions (1–5 dots). This yielded a five-component vector describing the neuron’s average response amplitude as a function of presented numerosity (Fig. 2c, Supplementary Fig. S7). The preferred numerosity of each neuron was defined as the numerosity corresponding to the maximum response in its tuning curve. To obtain single fish-level tuning profiles, tuning curves from all number-selective neurons for each fish were grouped according to their preferred numerosity and averaged, resulting in a single fish-averaged tuning curve for each preferred numerosity. These were further averaged across the multiple fish to produce the population averages for each age cohort (Fig. 2d).

To examine the overall numerical coding structure, averaging across all five types of preferred numerosity, we next aligned and averaged all age-group fish-averaged tuning curves according to their numerical distance from the preferred numerosity (Fig. 2e, Supplementary Fig. S8). In this representation, the x-axis denotes the numerical distance (from –4 to +4), where 0 corresponds to the preferred numerosity, –1 to one numerosity smaller (e.g., the response of number-3 neurons to 2 dots), and +1 to one numerosity larger (e.g., the response of number-3 neurons to 4 dots). The resulting curves, shown in Supplementary Fig. S8, capture how neuronal responses decrease with increasing numerical distance from the preferred numerosity. Finally, to quantify this effect irrespective of direction, responses were averaged by absolute numerical distance (e.g., responses to both 1 and 5 for a 3-tuned neuron correspond to a distance of 2), producing the population-level overall differential response shown in Fig. 2e.

### Brain spatial registration and region segmentation

All samples were first registered to a brain template of each respective age group with ITKsnap (40) using the average Ca^2+^ signal in time. All identified neuron centers were remapped to the final brain template to compare across different fish. To identify subregions of the forebrain, we registered the brain templates to the mapZebrain atlas (73) using affine transformation, then selected the available subregion Boolean masks.

### Support Vector Machine decoder

To assess the predictive capacity of the number-selective neurons, we implemented a supervised classification analysis based on a Support Vector Machine (SVM) with a linear kernel. This approach was chosen because (i) linear SVMs are widely used in numerosity decoding studies, facilitating comparison with previous work (45); (ii) they are robust to overfitting in the presence of trial-to-trial variability and moderate sample sizes (48 trials per numerosity condition in our case); and (iii) they do not assume normality or equal covariance across classes and are generally less sensitive to outliers than logistic regression. For each zebrafish, a separate decoder was constructed using the Ca² activity from each of the identified number neurons (that is, the full ensemble of number-selective neurons) as input features. Specifically, for each trial, we computed the mean ΔF/F activity of every number-selective neuron during a 3-s window encompassing the 1-s stimulus period and the subsequent 2-s post-stimulus period. These activities formed an n-dimensional input vector (where *n* = number of number-selective neurons from that fish), with each dimension corresponding to one neuron’s response, and the five stimulus numerosities (1–5 dots) serving as class labels.

Each SVM decoder was trained and tested within a leave-one-out (LOO) cross-validation framework. The model was trained on data from 7 of the 8 super-trials (Trainin*g* set) and evaluated on the remaining LOO super-trial (Testing set), iterating over all eight possible left-out supertrials. This yielded eight independent decoding results per fish, which were then averaged to obtain the single fish-level decoding performance. As a control for chance-level performance, we applied the same decoding procedure to the no-stimulus cohort, using all segmented neurons as features (since they lacked significant number-selective neurons), providing a matched control for potential non-numerical confounds. Confusion matrices showing the prediction performance for each single-fish decoder are shown in Supplementary Fig. S12. Finally, we averaged the performance across all samples within each age cohort to yield the results shown in Fig. 4b-c.

### Verification of stability of results across the recording time

To verify that the 90-minute imaging sessions did not introduce time-dependent confounding effects, we conducted a control analysis assessing the temporal stability of our results. Each 90-minute recording consisted of eight consecutive supertrials, each lasting 11.25 minutes. This temporal structure allowed us to test whether the outcomes of our analyses varied systematically over time. We leveraged the existing LOO framework to evaluate stability across the recording. Since each LOO round excluded a different supertrial, we could directly examine whether the results depended on the temporal position of the left-out supertrial (from position 1 at the beginning to position 8 at the end of the recording). For each LOO round, we quantified (i) the identity of the number-selective neurons identified by the permutation ANOVA and (ii) the prediction accuracy of the SVM decoder. These metrics were then plotted as a function of the supertrial order to assess potential temporal trends in neuron selection or decoding performance (Supplementary Fig. S6).

### Statistical analysis

Statistical analyses and graph preparation were conducted using custom Python scripts, Seaborn library (74), and Inkscape.

## Author contributions

Conceptualization: CHB, GV, SEF, PL, AN, TVT

Setup implementation: PL, AN, KKD, MJ, TVT

Methodology: PL, AN, MZ, TVT

Investigation: PL, AN, MZ, NM, KKD, MJ, AM, MEMP, JVTP, CHB, GV, SEF, TVT

Code writing: PL, AN, MZ

Visualization: PL, AN, MZ, SEF, TVT

Funding acquisition: KKD, CHB, GV, SEF, TVT

Supervision: CHB, GV, SEF, TVT

Writing – original draft: PL, AN, MZ, SEF, TVT

Writing – review & editing: PL, MZ, AN, AM, MEMP, JVTP, KKD, CHB, GV, SEF, TVT

## Classification

Biological Sciences, Neuroscience

## Acknowledgements

We thank David Prober (Caltech) for sharing the zebrafish used in this study and Falk Schneider for assistance with editing the manuscript.

## Funding

Human Frontier Science Program Grant RGP0008/2018 to CHB, SEF, GV.

ERC European Union’s Horizon 2020 Research and Innovation Program Grant Agreement 833504 – SPANUMBRA to GV, CHB.

FARE–Ricerca in Italia: Framework per l’Attrazione ed il Rafforzamento delle Eccellenze per la ricerca in Italia, III edizione, project “NUMBRISH – The neurobiology of numerical cognition: searching for a molecular signature in the zebrafish brain” Prot. R20YL9WN9N to GV.

PRIN—Progetti di rilevante Interesse Nazionale 2022—PNRR (Grant Agreement P2022TKY7B: “The emergence of proto-arithmetic abilities with empty and non-empty sets” to GV.

National Institutes of Health 1U01NS122082-01, 1R34NS126800-01 to TVT.

Alred E. Mann Doctoral Fellowship to KKD.

## Data and materials availability

Data, code and materials used in the analysis are available at: https://doi.org/10.6084/m9.figshare.27676197.v3. Additional links to the tools used are available in Supplementary Table S10.

## References

1. Macchinizzi M, Felisatti A, Rugani R. The Social Relevance of Numbers: Insights from Animal Studies. Life. 2025 Nov;15(11):1775.

2. Viswanathan P, Stein AM, Nieder A. Sequential neuronal processing of number values, abstract decision, and action in the primate prefrontal cortex. PLOS Biology. 2024 Feb 16;22(2):e3002520.

3. Liang T, Peng RC, Rong KL, Li JX, Ke Y, Yung WH. Disparate processing of numerosity and associated continuous magnitudes in rats. Science Advances [Internet]. 2024 Feb 23 [cited 2026 Jan 7]; Available from: https://www.science.org/doi/10.1126/sciadv.adj2566

4. Bengochea M, Sitt JD, Izard V, Preat T, Cohen L, Hassan BA. Numerical discrimination in Drosophila melanogaster. Cell Reports [Internet]. 2023 July 25 [cited 2026 Jan 7];42(7). Available from: https://www.cell.com/cell-reports/abstract/S2211-1247(23)00783-0

5. Cross FR, Jackson RR. Representation of different exact numbers of prey by a spider-eating predator. Interface Focus. 2017 Apr 21;7(3):20160035.

6. C. N. Templeton, E. Greene, Nuthatches eavesdrop on variations in heterospecific chickadee mobbing alarm calls. Proc. Natl. Acad. Sci. U. S. A. 104, 5479–5482 (2007).

7. Edwards CJ, Alder TB, Rose GJ. Auditory midbrain neurons that count. Nat Neurosci. 2002 Oct;5(10):934–6.

8. Alder TB, Rose GJ. Integration and recovery processes contribute to the temporal selectivity of neurons in the midbrain of the northern leopard frog, Rana pipiens. J Comp Physiol A. 2000 Oct 1;186(10):923–37.

9. Brannon EM, Merritt DJ. Evolutionary foundations of the approximate number system. In: Space, time and number in the brain: Searching for the foundations of mathematical thought. San Diego, CA, US: Elsevier Academic Press; 2011. p. 207–24.

10. Piazza M. Neurocognitive start-up tools for symbolic number representations. Trends in Cognitive Sciences. 2010 Dec 1;14(12):542–51.

11. Piazza M, Izard V, Pinel P, Le Bihan D, Dehaene S. Tuning Curves for Approximate Numerosity in the Human Intraparietal Sulcus. Neuron. 2004 Oct 28;44(3):547–55.

12. Xu F, Spelke ES. Large number discrimination in 6-month-old infants. Cognition. 2000 Jan 10;74(1):B1–11.

13. Ditz HM, Nieder A. Neurons selective to the number of visual items in the corvid songbird endbrain. Proceedings of the National Academy of Sciences. 2015 June 23;112(25):7827–32.

14. Ditz HM, Nieder A. Sensory and Working Memory Representations of Small and Large Numerosities in the Crow Endbrain. J Neurosci. 2016 Nov 23;36(47):12044–52.

15. Messina A, Potrich D, Schiona I, Sovrano VA, Fraser SE, Brennan CH, et al. Neurons in the Dorso-Central Division of Zebrafish Pallium Respond to Change in Visual Numerosity. Cereb Cortex. 2022 Jan 15;32(2):418–28.

16. Messina A, Potrich D, Perrino M, Sheardown E, Miletto Petrazzini ME, Luu P, et al. Quantity as a Fish Views It: Behavior and Neurobiology. Frontiers in Neuroanatomy [Internet]. 2022 [cited 2023 Oct 31];16. Available from: https://www.frontiersin.org/articles/10.3389/fnana.2022.943504

17. Pfeifer JH, Allen NB, Byrne ML, Mills KL. Modeling Developmental Change: Contemporary Approaches to Key Methodological Challenges in Developmental Neuroimaging. Developmental Cognitive Neuroscience. 2018 Oct 1;33:1–4.

18. Viswanathan P, Nieder A. Neuronal correlates of a visual “sense of number” in primate parietal and prefrontal cortices. Proceedings of the National Academy of Sciences. 2013 July 2;110(27):11187–92.

19. Collins E, Park J, Behrmann M. Numerosity representation is encoded in human subcortex. Proceedings of the National Academy of Sciences. 2017 Apr 4;114(14):E2806–15.

20. Georgy L, Celeghin A, Marzi CA, Tamietto M, Ptito A. The superior colliculus is sensitive to gestalt-like stimulus configuration in hemispherectomy patients. Cortex. 2016 Aug 1;81:151–61.

21. Lorenzi E, Perrino M, Messina A, Zanon M, Vallortigara G. Innate responses to numerousness reveal neural activation in different brain regions in newly-hatched visually naïve chicks. Heliyon. 2024 July 30;10(14):e34162.

22. Eles JR, Vazquez AL, Kozai TDY, Cui XT. *In vivo* imaging of neuronal calcium during electrode implantation: Spatial and temporal mapping of damage and recovery. Biomaterials. 2018 Aug 1;174:79–94.

23. Ferguson M, Sharma D, Ross D, Zhao F. A Critical Review of Microelectrode Arrays and Strategies for Improving Neural Interfaces. Advanced Healthcare Materials. 2019;8(19):1900558.

24. Truong TV, Supatto W, Koos DS, Choi JM, Fraser SE. Deep and fast live imaging with two-photon scanned light-sheet microscopy. Nat Methods. 2011 Sept;8(9):757–60.

25. Wolf S, Supatto W, Debrégeas G, Mahou P, Kruglik SG, Sintes JM, et al. Whole-brain functional imaging with two-photon light-sheet microscopy. Nature Methods. 2015 May;12(5):379–80.

26. Keomanee-Dizon K, Fraser SE, Truong TV. A versatile, multi-laser twin-microscope system for light-sheet imaging. Review of Scientific Instruments. 2020 May 1;91(5):053703.

27. Vito G de, Turrini L, Müllenbroich C, Ricci P, Sancataldo G, Mazzamuto G, et al. Fast whole-brain imaging of seizures in zebrafish larvae by two-photon light-sheet microscopy. Biomed Opt Express, BOE. 2022 Mar 1;13(3):1516–36.

28. Nadtochiy A, Luu P, Fraser SE, Truong TV. VoDEx: a Python library for time annotation and management of volumetric functional imaging data. Bioinformatics. 2023 Sept 12;btad568.

29. Ahrens MB, Orger MB, Robson DN, Li JM, Keller PJ. Whole-brain functional imaging at cellular resolution using light-sheet microscopy. Nature Methods. 2013 May;10(5):413–20.

30. Choi TY, Choi TI, Lee YR, Choe SK, Kim CH. Zebrafish as an animal model for biomedical research. Exp Mol Med. 2021 Mar;53(3):310–7.

31. Doszyn O, Dulski T, Zmorzynska J. Diving into the zebrafish brain: exploring neuroscience frontiers with genetic tools, imaging techniques, and behavioral insights. Front Mol Neurosci. 2024 Mar 12;17.

32. Dana H, Sun Y, Mohar B, Hulse BK, Kerlin AM, Hasseman JP, et al. High-performance calcium sensors for imaging activity in neuronal populations and microcompartments. Nat Methods. 2019 July;16(7):649–57.

33. Adam E, Zanon M, Messina A, Vallortigara G. Looks like home: numerosity, but not spatial frequency guides preference in zebrafish larvae (Danio rerio). Anim Cogn. 2024;27(1):53.

34. Lucon-Xiccato T, Gatto E, Fontana CM, Bisazza A. Quantity discrimination in newly hatched zebrafish suggests hardwired numerical abilities. Commun Biol. 2023 Mar 23;6(1):247.

35. Sheardown E, Torres-Perez JV, Anagianni S, Fraser SE, Vallortigara G, Butterworth B, et al. Characterizing ontogeny of quantity discrimination in zebrafish. Proc Biol Sci. 2022 Feb 9;289(1968):20212544.

36. Lorenzi E, Perrino M, Messina A, Zanon M, Vallortigara G. A Kaspar Hauser experiment for innateness of numerical cognition. bioRxiv; 2023 [cited 2026 Jan 15]. p. 2023.12.06.570352.

37. Zanon M, Potrich D, Bortot M, Vallortigara G. Towards a standardization of non-symbolic numerical experiments: GeNEsIS, a flexible and user-friendly tool to generate controlled stimuli. Behav Res. 2022 Feb 1;54(1):146–57.

38. Haug MF, Biehlmaier O, Mueller KP, Neuhauss SC. Visual acuity in larval zebrafish: behavior and histology. Front Zool. 2010 Mar 1;7(1):8.

39. Avants B, Tustison NJ, Song G. Advanced Normalization Tools: V1.0. The Insight Journal [Internet]. 2009 July 29 [cited 2026 Jan 20]; Available from: https://www.insight-journal.org/browse/publication/681

40. Yushkevich PA, Piven J, Hazlett HC, Smith RG, Ho S, Gee JC, et al. User-guided 3D active contour segmentation of anatomical structures: Significantly improved efficiency and reliability. NeuroImage. 2006 July 1;31(3):1116–28.

41. Giovannucci A, Friedrich J, Gunn P, Kalfon J, Brown BL, Koay SA, et al. CaImAn an open source tool for scalable calcium imaging data analysis. Kleinfeld D, King AJ, editors. eLife. 2019 Jan 17;8:e38173.

42. Yang E, Zwart MF, James B, Rubinov M, Wei Z, Narayan S, et al. A brainstem integrator for self-location memory and positional homeostasis in zebrafish. Cell. 2022 Dec 22;185(26):5011–5027.e20.

43. Li Q, Uitto J. Zebrafish as a Model System to Study Skin Biology and Pathology. Journal of Investigative Dermatology. 2014 June 1;134(6):1–6.

44. Kimmel CB, Ballard WW, Kimmel SR, Ullmann B, Schilling TF. Stages of embryonic development of the zebrafish. Developmental Dynamics. 1995;203(3):253–310.

45. Kirschhock ME, Nieder A. Number selective sensorimotor neurons in the crow translate perceived numerosity into number of actions. Nat Commun. 2022 Nov 14;13(1):6913.

46. Luu P, Fraser SE, Schneider F. More than double the fun with two-photon excitation microscopy. Commun Biol. 2024 Mar 26;7(1):364.

47. Kobylkov D, Mayer U, Zanon M, Vallortigara G. Number neurons in the nidopallium of young domestic chicks. Proc Natl Acad Sci U S A. 2022 Aug 9;119(32):e2201039119.

48. Izard V, Sann C, Spelke ES, Streri A. Newborn infants perceive abstract numbers. Proceedings of the National Academy of Sciences. 2009 June 23;106(25):10382–5.

49. Easter Jr Stephen S, Nicola GN. The Development of Vision in the Zebrafish (*Danio rerio*). Developmental Biology. 1996 Dec 15;180(2):646–63.

50. Borla MA, Palecek B, Budick S, O’Malley DM. Prey Capture by Larval Zebrafish: Evidence for Fine Axial Motor Control. Brain Behav Evol. 2002 Dec 2;60(4):207–29.

51. Nieder A. The neural code for number. In: Space, time and number in the brain: Searching for the foundations of mathematical thought. San Diego, CA, US: Elsevier Academic Press; 2011. p. 103–18.

52. Nieder A, Freedman DJ, Miller EK. Representation of the Quantity of Visual Items in the Primate Prefrontal Cortex. Science. 2002 Sept 6;297(5587):1708–11.

53. Park HC, Hong SK, Kim HS, Kim SH, Yoon EJ, Kim CH, et al. Structural comparison of zebrafish Elav/Hu and their differential expressions during neurogenesis. Neuroscience Letters. 2000 Jan 28;279(2):81–4.

54. Tabor KM, Marquart GD, Hurt C, Smith TS, Geoca AK, Bhandiwad AA, et al. Brain-wide cellular resolution imaging of Cre transgenic zebrafish lines for functional circuit-mapping. Raman IM, Stainier DY, Wyart C, editors. eLife. 2019 Feb 8;8:e42687.

55. Cheng RK, Jesuthasan SJ, Penney TB. Zebrafish forebrain and temporal conditioning. Philos Trans R Soc Lond B Biol Sci. 2014 Mar 5;369(1637):20120462.

56. Medina L, Abellán A, Desfilis E. Evolution of Pallial Areas and Networks Involved in Sociality: Comparison Between Mammals and Sauropsids. Front Physiol. 2019 July 12 ;10.

57. Förster D, Helmbrecht TO, Mearns DS, Jordan L, Mokayes N, Baier H. Retinotectal circuitry of larval zebrafish is adapted to detection and pursuit of prey. Kawakami K, King AJ, Kawakami K, Du J, editors. eLife. 2020 Oct 12;9:e58596.

58. Gazzola A, Vallortigara G, Pellitteri-Rosa D. Continuous and discrete quantity discrimination in tortoises. Biol Lett. 2018 Dec 12;14(12):20180649.

59. Muto A, Ohkura M, Abe G, Nakai J, Kawakami K. Real-time visualization of neuronal activity during perception. Curr Biol. 2013 Feb 18;23(4):307–11.

60. Bengochea M, Hassan B. Numerosity as a visual property: Evidence from two highly evolutionary distant species. Front Physiol. 2023 Feb 9;14.

61. Lorenzi E, Perrino M, Vallortigara G. Numerosities and Other Magnitudes in the Brains: A Comparative View. Front Psychol. 2021 Apr 15;12.

62. Ritchie JB, Kaplan DM, Klein C. Decoding the Brain: Neural Representation and the Limits of Multivariate Pattern Analysis in Cognitive Neuroscience. The British Journal for the Philosophy of Science. 2019 June;70(2):581–607.

63. Moyer AJ, Chrabasz JA, Barcus AK, Cheng J, Capps MES, Lalonde RL, et al. Genetic context of transgene insertion can influence neurodevelopment in zebrafish. Genetics. 2025 Nov 1;231(3):iyaf195.

64. Zhang Y, Rózsa M, Liang Y, Bushey D, Wei Z, Zheng J, et al. Fast and sensitive GCaMP calcium indicators for imaging neural populations. Nature. 2023 Mar;615(7954):884–91.

65. Avdesh A, Chen M, Martin-Iverson MT, Mondal A, Ong D, Rainey-Smith S, et al. Regular Care and Maintenance of a Zebrafish (Danio rerio) Laboratory: An Introduction. Journal of Visualized Experiments (JoVE). 2012 Nov 18;(69):e4196.

66. Vito G de, Vito G de, Ricci P, Turrini L, Turrini L, Gavryusev V, et al. Effects of excitation light polarization on fluorescence emission in two-photon light-sheet microscopy. Biomed Opt Express, BOE. 2020 Aug 1;11(8):4651–65.

67. Edelstein A, Amodaj N, Hoover K, Vale R, Stuurman N. Computer Control of Microscopes Using µManager. Current Protocols in Molecular Biology. 2010;92(1):14.20.1-14.20.17.

68. Temizer I, Donovan JC, Baier H, Semmelhack JL. A Visual Pathway for Looming-Evoked Escape in Larval Zebrafish. Current Biology. 2015 July 20;25(14):1823–34.

69. Peirce J, Gray JR, Simpson S, MacAskill M, Höchenberger R, Sogo H, et al. PsychoPy2: Experiments in behavior made easy. Behav Res. 2019 Feb 1;51(1):195–203.

70. Bahl A, Engert F. Neural circuits for evidence accumulation and decision making in larval zebrafish. Nat Neurosci. 2020 Jan;23(1):94–102.

71. Koning HK, Ahemaiti A, Boije H. A deep-dive into fictive locomotion – a strategy to probe cellular activity during speed transitions in fictively swimming zebrafish larvae. Biol Open. 2022 Mar 22;11(3):bio059167.

72. Field A, Miles J, Field Z. Chapter 12: Factorial ANOVA. In: Discovering statistics using R. SAGE Publications Ltd STM; 2012. p. 498–546.

73. Kunst M, Laurell E, Mokayes N, Kramer A, Kubo F, Fernandes AM, et al. A Cellular-Resolution Atlas of the Larval Zebrafish Brain. Neuron. 2019 July 3;103(1):21–38.e5.

74. Waskom ML. seaborn: statistical data visualization. Journal of Open Source Software. 2021 Apr 6;6(60):3021.

