## Supplementary materials for "Ordered developmental emergence of number-selective neurons in days-old zebrafish"

### Supplementary Materials
#### for
#### *Ordered developmental emergence of number-selective neurons in days-old zebrafish*

**Authors:**

Peter Luu [*], Anna Nadtochiy [*], Mirko Zanon[*], Noah Moreno, Andrea Messina, Maria Elena Miletto Petrazzini, Jose Vicente Torres Perez, Kevin Keomanee-Dizon, Matthew Jones, Caroline H. Brennan, Giorgio Vallortigara, Scott E. Fraser, Thai V. Truong[#]

**Content:**

Supplementary Figures S1-S12
Supplementary Tables S1-S10
Supplemental Results

**Supplementary Fig. S1. Sequence of stimuli including all possible combinations of spread and sizes using a new pattern for each stimulus.**

Each stimulus lasts for 1 second, and the inter-stimulus duration varies between 15 and 27 seconds. A stimulus cycle is 684 seconds in total when including both convex hull and inter-distance controls. When a pseudo-random cycle is repeated, a novel dot pattern is displayed. The cycle is repeated 8 times per sample.

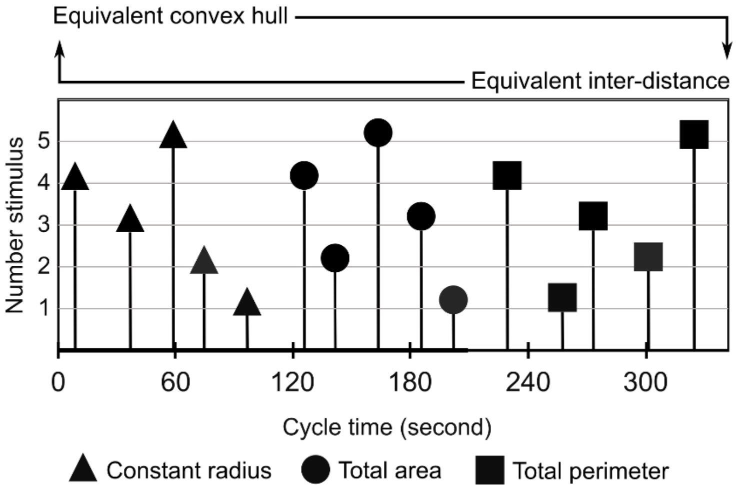

**Supplementary Fig. S2. Example of segmentation output using the CaImAn toolbox.**

Image panels show the neuronal segmentation results obtained with the CaImAn toolbox. Segmented neurons are indicated by black circles outlined in white, overlaid on the corresponding $Ca^{2+}$ fluorescence images, with fluorescence intensity color-coded according to the look-up table shown at the bottom. Eight representative z-slices are displayed, each taken from a different depth within the zebrafish brain (z = 0 corresponds to the dorsal surface; rostral is to the left).

Segmentation was performed on a restricted temporal window (3 seconds before and 5 seconds after each stimulus presentation) to emphasize neurons responsive to visual stimuli. As a result, some neurons that were well captured in the imaging field but not visually-responsive might not appear segmented. This approach biases the segmentation toward functionally active neurons, which is desirable for subsequent analyses of numerosity selectivity.

Following segmentation, we applied stringent filtering criteria based on signal-to-noise ratio and stimulus-driven activity to exclude spurious or noisy components (see Methods for details). The example images shown here illustrate the raw segmentation output; only a subset of these units passed further statistical screening (permutation ANOVA) and were classified as number-selective neurons.

**Supplementary Fig. S2** (see caption on previous page).

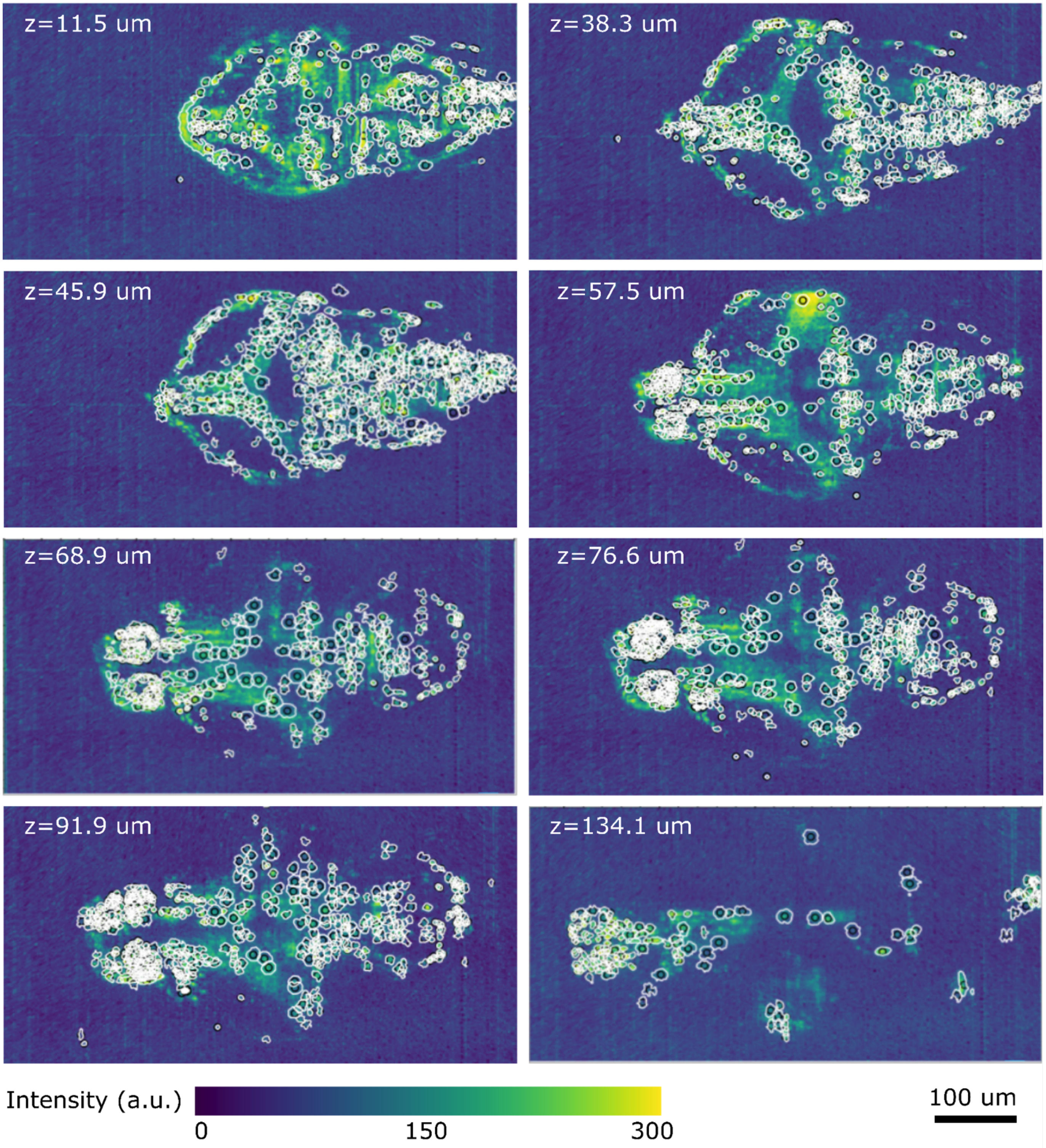

**Supplementary Fig. S3. Example of $Ca^{2+}$ signal traces from a number-selective neuron, illustrating responsiveness to changes in numerosity rather than to geometrical covariates.**

The normalized $Ca^{2+}$ activity is shown for the peristimulus window (3 s before, 1 s stimulus, 5 s after), averaged across multiple trials. The trials are grouped, averaged, and presented according to the five geometrical covariates that are kept identical across different trials (see Methods): radius (a), total area (b), total perimeter (c), total convex hull (d), and inter-distance (e). In each panel, the gray bars mark the time window of the different stimuli (numerosity 1 to 5 dots, from left to right), tick marks on the horizontal axis represent seconds, and error bars denote the SEM across trials. Red lines indicate the mean $Ca^{2+}$ response across all experimental trials, while black lines indicate the mean response for trials where the numerical stimuli were controlled for a specific covariate.

For example, in (a), trials where the radius of the dots are kept constant are averaged for each numerical stimulus (black curves). These curves display the same behavior as the ones constructed by taking into account the entire set of trials (red curves). Analogous results can be found for total area (b), total perimeter (c), convex hull (d) and inter-distance (e).

This example visually demonstrates how the permutation ANOVA procedure (see Methods) identifies neurons whose responses are selective for numerosity and not explained by non-numerical covariates. In practice, the ANOVA includes both numerosity and control conditions (total area, total perimeter, radius, convex hull, inter-distance) as factors. Neurons are considered number-selective when they show a significant effect of numerosity while not showing significant modulation by any of the control conditions. By applying this criterion, the procedure ensures that the observed responses cannot be attributed to variations in the non-numerical stimulus dimensions, but instead reflect selectivity for numerosity per se.

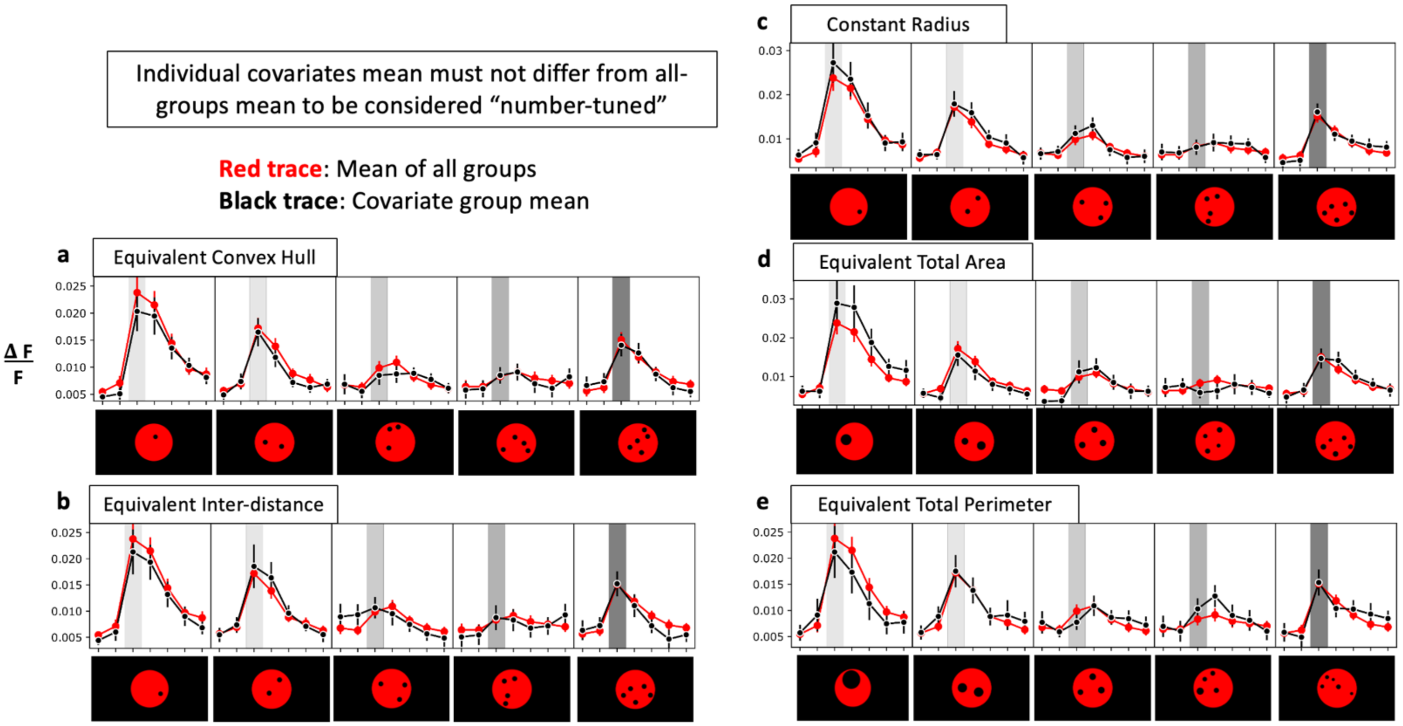

**Supplementary Fig. S4. High degree of overlap in number neurons found from different Analysis groups of the experimental data.**

See Methods for details. The overlap percentage of identified number neurons, for all possible 8 pair-wise comparisons, is plotted as a color-coded heat map, ranging from 0 to 100%. The no-stimulus cohort should yield no significant number-selective neurons and thus provides a baseline representing false positives. The matrices are symmetrical with each Analysis group represented as an entry across both horizontal and vertical axes. The overlap percentages are consistently higher than chance-level of 50% for all the experimental results across the 3 age cohorts ($p < 1.05E-32$, one-side one-sample t-test, for the average overlap ratios, for each of the age cohort), while consistently at chance level for the no-stimulus control cohort ($p = 1.00$, one-side one-sample t-test, for the average overlap ratios).

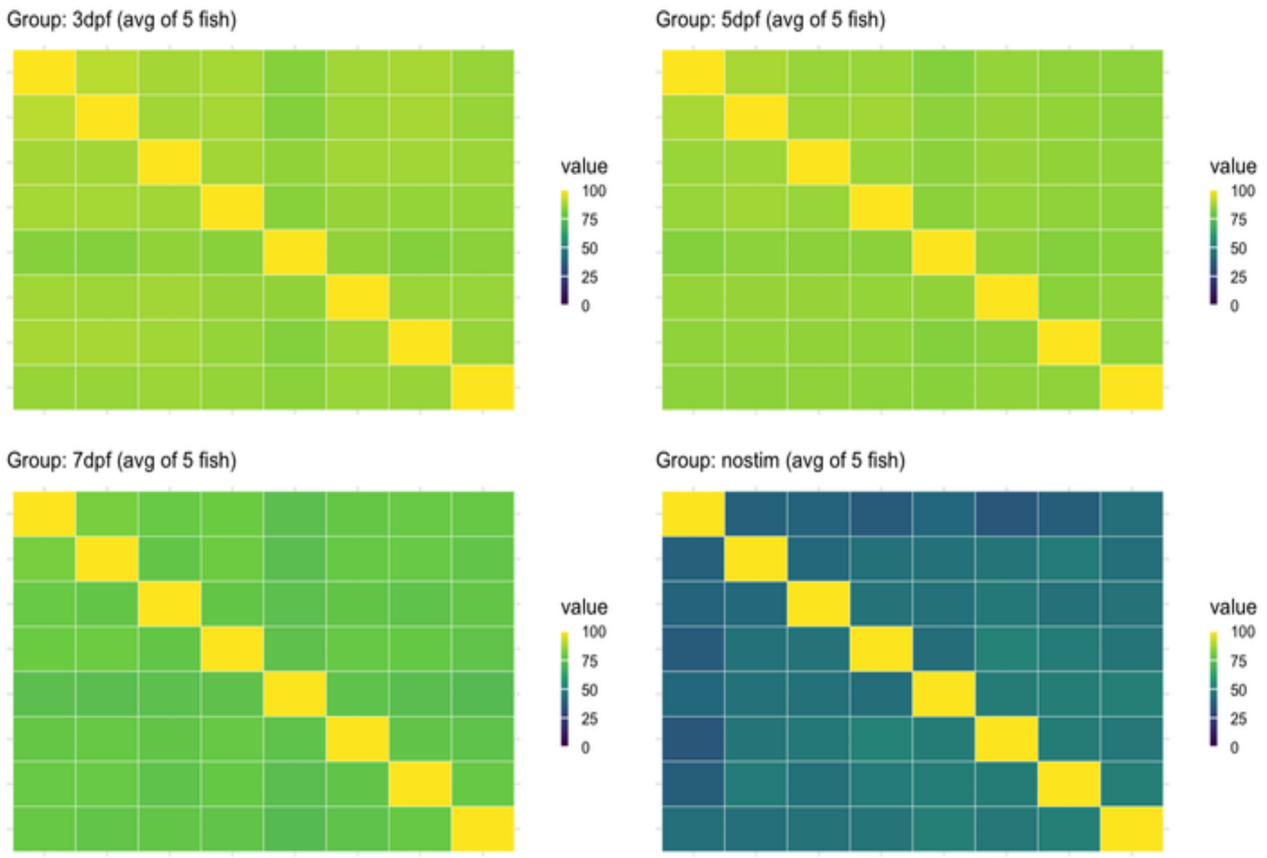

**Supplementary Fig. S5. High Tuning Correlation scores of the number neurons' tuning curves, compared between Analysis and Testing groups.**

See Methods for details. The no-stimulus cohort should yield no significant number-selective neurons and thus provides a baseline representing false positives. Shuffled datasets were generated by randomizing the numerosity labels of the corresponding real datasets and provide a baseline for chance. Statistical comparison with t-testing, yielding p-values of 0.00012, 0.0098, 0.0026, and 0.49, for the 3 dpf, 5 dpf, 7 dpf, and no-stimulus cohorts, respectively.

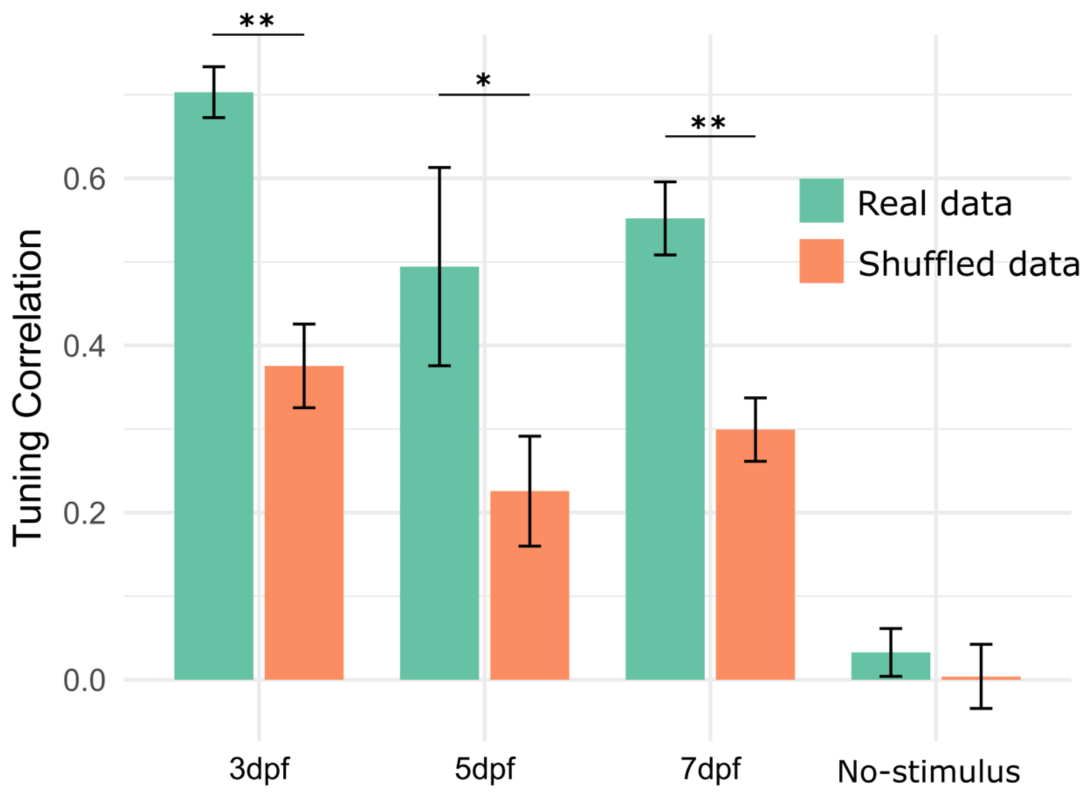

**Supplementary Fig. S6. Stability of number-selective neurons and decoding performance as a function of experimental recording time.**

Since the supertrials are arranged in temporal sequence, we examined analysis results as the leave-one-out (LOO) supertrial varies from 1 (beginning of recording) to 8 (end of recording), to assess whether there is any time-dependent effect from the 90-minute recording.

**(a)** Neuron identity overlap. For each fish, number-selective neurons were identified using a two-way permutation ANOVA (factors: Numerosity × Control condition) in a leave-one-out (LOO) cross-validation scheme. At each LOO fold, neurons were detected from seven training supertrials, and the percentage of overlapping neurons with the other LOO subsets was computed and averaged, yielding a single overlap value. Plotted are group means (± SEM) across fish within each age cohort (color-coded). This metric quantifies the reliability with which the same number-selective neurons are identified across different subsets of trials. Single LOO overlap values were compared to the group average across all supertrials (shown as column plot to the right; 3 dpf: 86.4%; 5 dpf: 85.4%; 7 dpf: 77.5%; No-stimulus: 44.4%) using one-sample t-tests, showing no significant differences across rounds.

**(b)** SVM decoding accuracy. To assess generalization performance, a linear support vector machine (SVM) was trained on the activity of number-selective neurons from seven supertrials and tested on the held-out supertrial. Classification accuracy for each fold was calculated as the proportion of correctly predicted trials (average diagonal of the confusion matrix). Accuracy values were averaged across fish within each developmental group (color-coded) to produce the curves (± SEM). The horizontal dotted line indicates chance-level accuracy (20%). Single LOO supertrial accuracies were compared to the group average across all supertrials (shown a column plot to the right; 3 dpf: 0.42; 5 dpf: 0.44; 7 dpf: 0.41; No-stimulus: 0.21), showing no significant differences across rounds (except for 5dpf at supertrial #3: $t=-3.4$, $p=0.03$).

**Supplementary Fig. S6** (see caption on previous page).

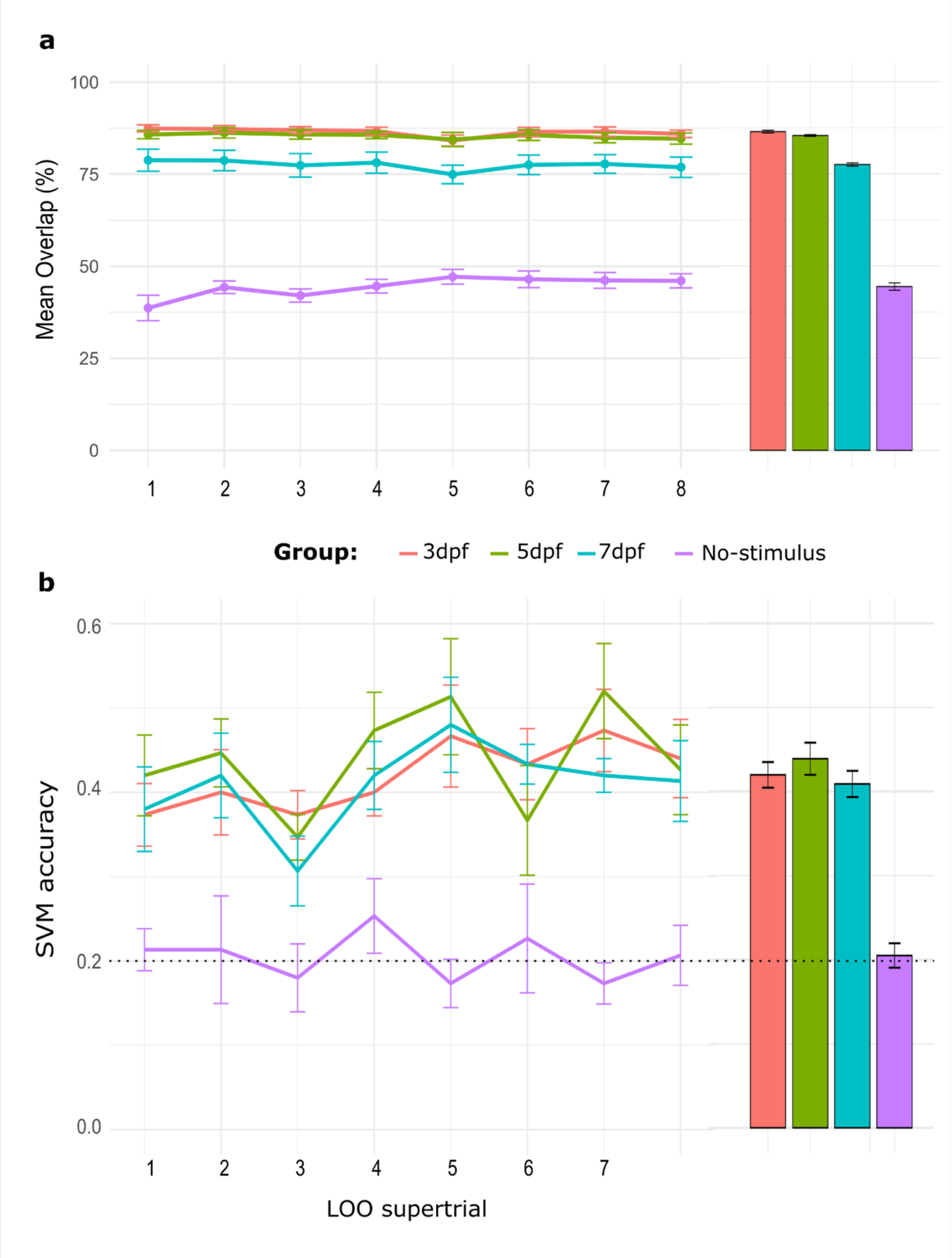

**Supplementary Fig. S7. Representative single-neuron numerosity tuning curves.**

Representative tuning curves are shown for each age cohort (a) 3 dpf, (b) 5 dpf, and (c) 7 dpf. Each raw correspond to a different fish, each column (color-coded) represents the average response of all the neurons preferring 1 to 5 dots. See Methods for how the tuning curve for each number-selective neuron was constructed.

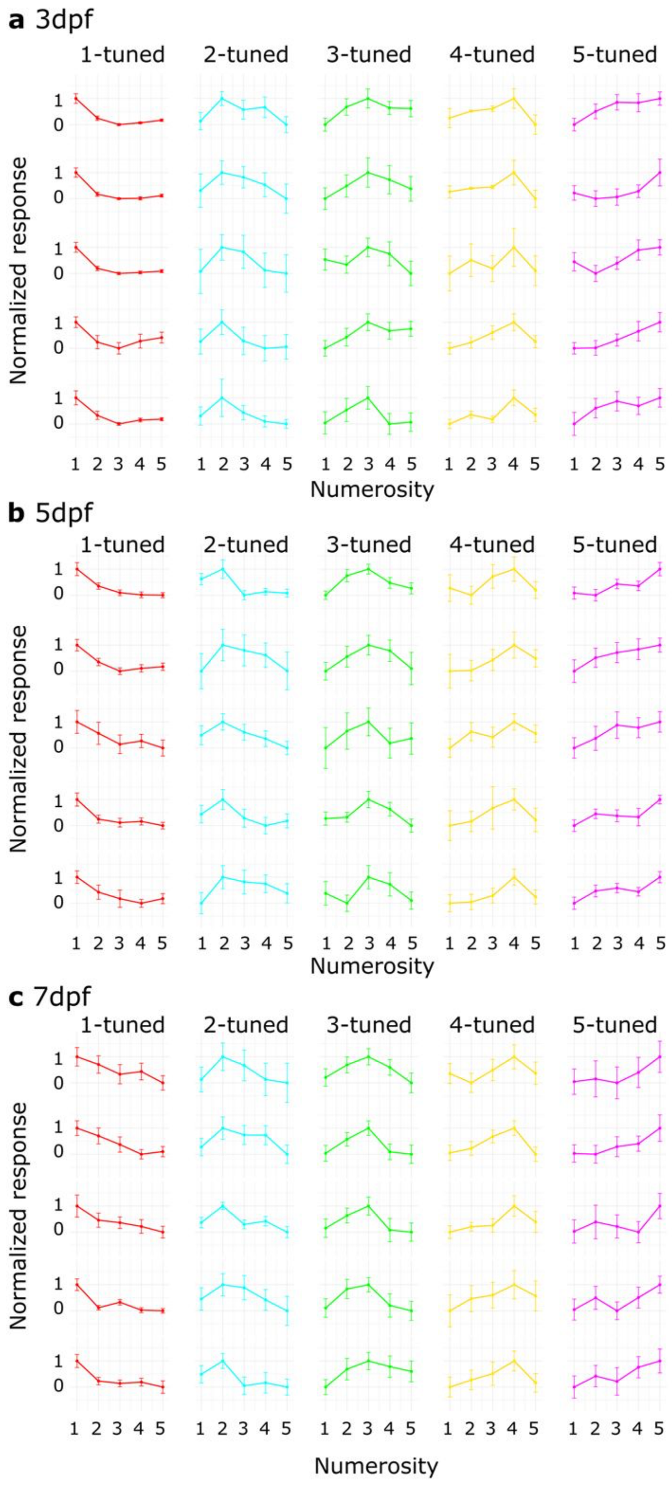

**Supplementary Fig. S8. Differential tuning response of 3, 5, and 7 dpf age groups.**

The overall differential tuning responses (analogous to those shown Fig. 2e), averaged across five fish per age group, represent the normalized average Ca2+ activity of all number-selective neurons in response to stimuli as functions of actual (signed) numerical distances from the preferred numerosity. The response of a neuron to its preferred numerosity is represented by 0.

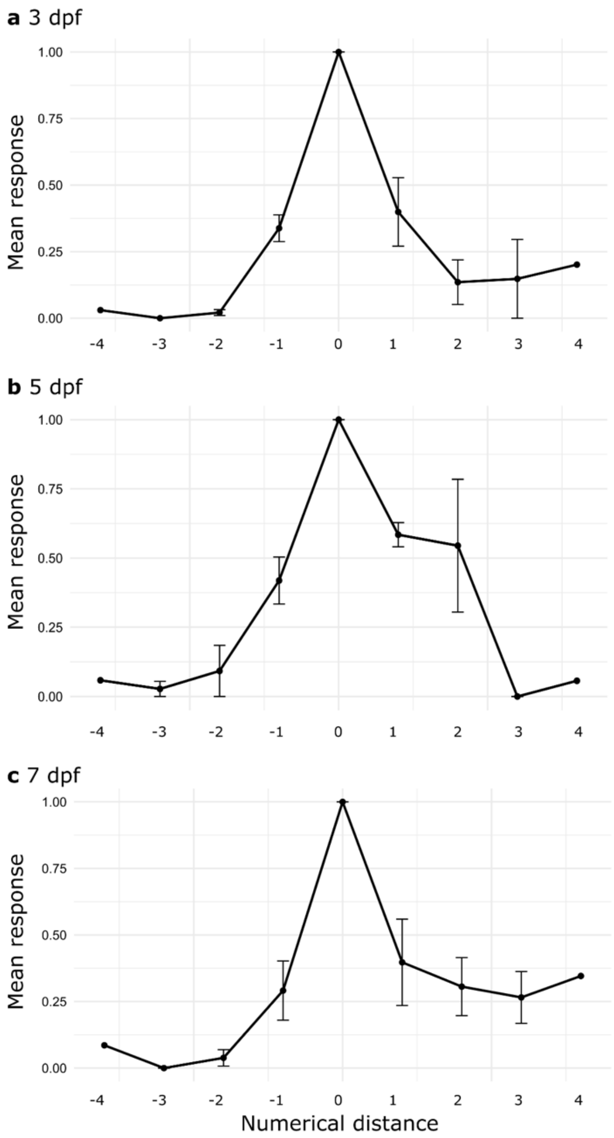

**Supplementary Fig. S9. No-stimulus negative control.**

Locations of number-selective neurons in one larval zebrafish as orthographic projections without presenting number stimuli. The white circles represent the centers of each identified number-selective neuron. Scale bar: 100 μm.

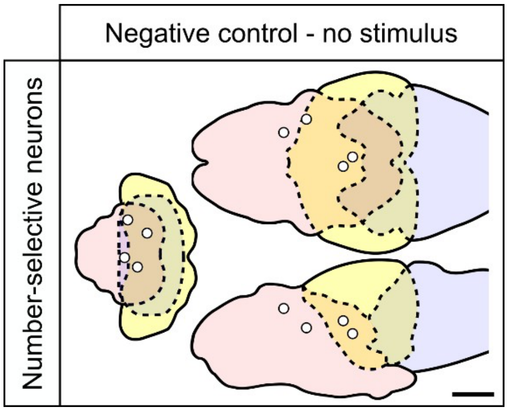

**Supplementary Fig. S10. Localization of number-selective neurons at three stages of development.**

**a** The 3D map of the brain was divided into three major brain regions (forebrain, midbrain, hindbrain). Solid lines indicate delineation of major brain regions; dash lines indicate overlapping regions.

**b** Locations of number-selective neurons in three different individual larval zebrafish at three stages of development, representing the results as point maps in orthographic projections. The individual-colored dots represent the centers of each number-selective neuron. Columns indicate age; neurons responding with specific number tunings are shown as rows. Scale bar: 100 µm.

**Supplementary Fig. S10** (see caption on previous page).

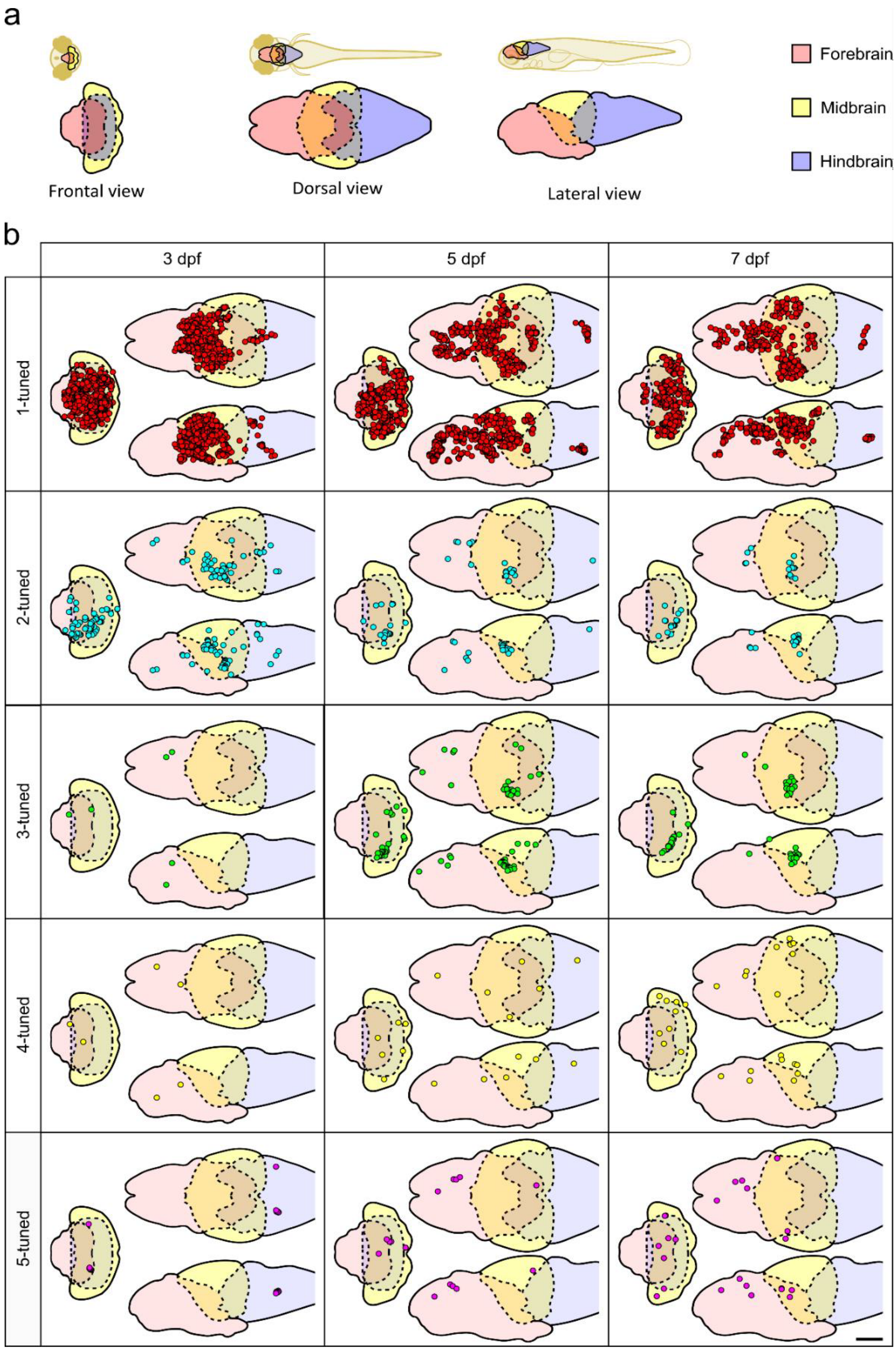

**Supplementary Fig. S11. Distribution of sub-regional number-selective neurons normalized by total number-selective neurons in the forebrain.**

See Table S6 for p-values and f-scores.

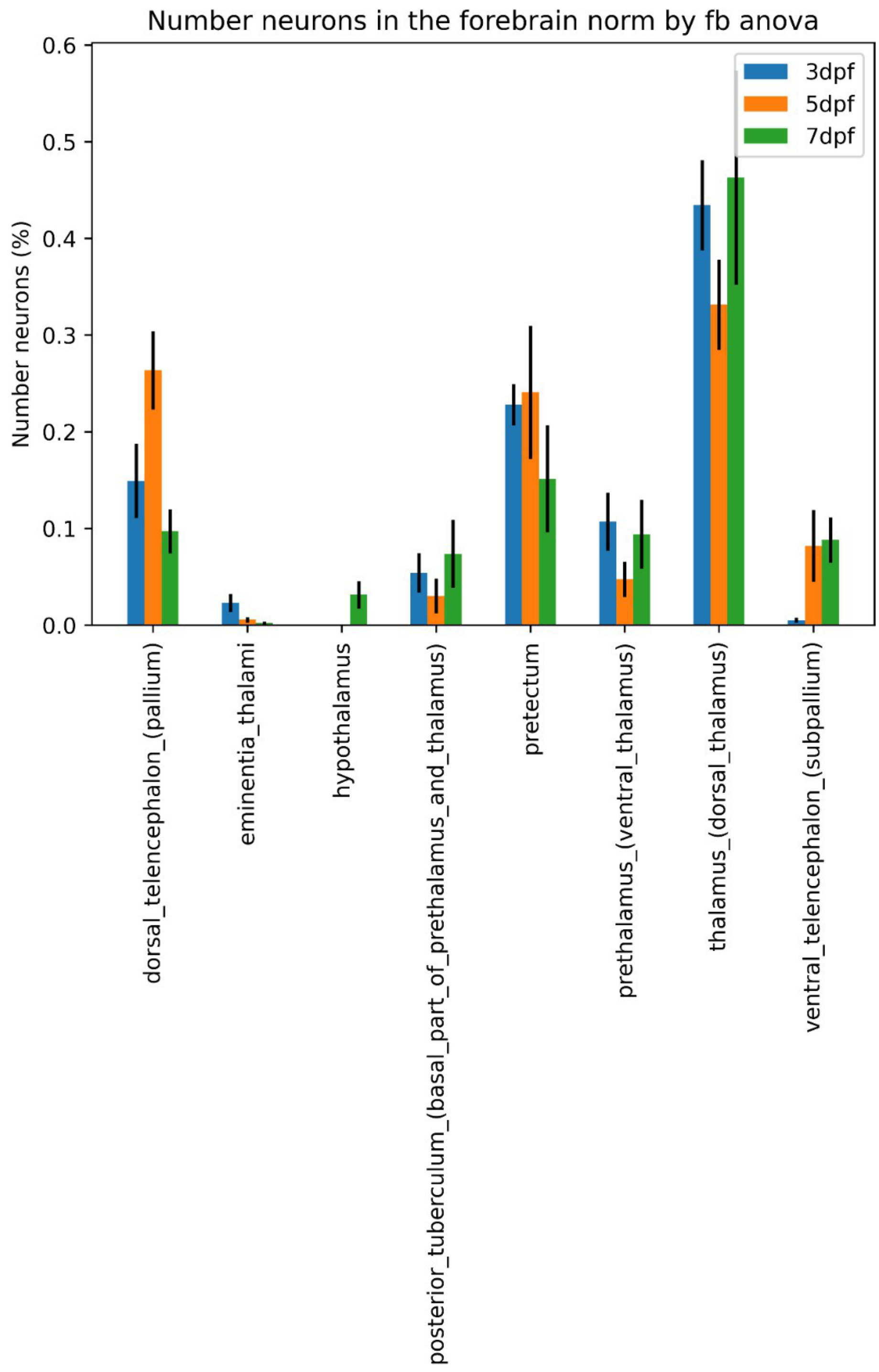

**Supplementary Fig. S12. Performance of the SVM decoder in predicting the stimulus presentation, for individual zebrafish samples.**

Confusion matrix and mean accuracy of the support vector machine (SVM) single-fish decoder (see Methods). Matrix entries correspond to percentages, defined as the ratio of the number of correct predictions over the total instances of each true label. The mean values plotted in the column graph are averages of the diagonal terms in the corresponding matrix. Each matrix and column is labeled with the ID of the corresponding fish. Random chance = 20% (1 out of 5 choices).

All decoders show high predictive power, significantly above what would be expected from chance, with the mean accuracies exceeding chance probability by more than 2-fold. The associated means and p-values for the statistical significance tests are listed in Supplementary Table S10.

**Supplementary Fig. S12** (see caption on previous page).

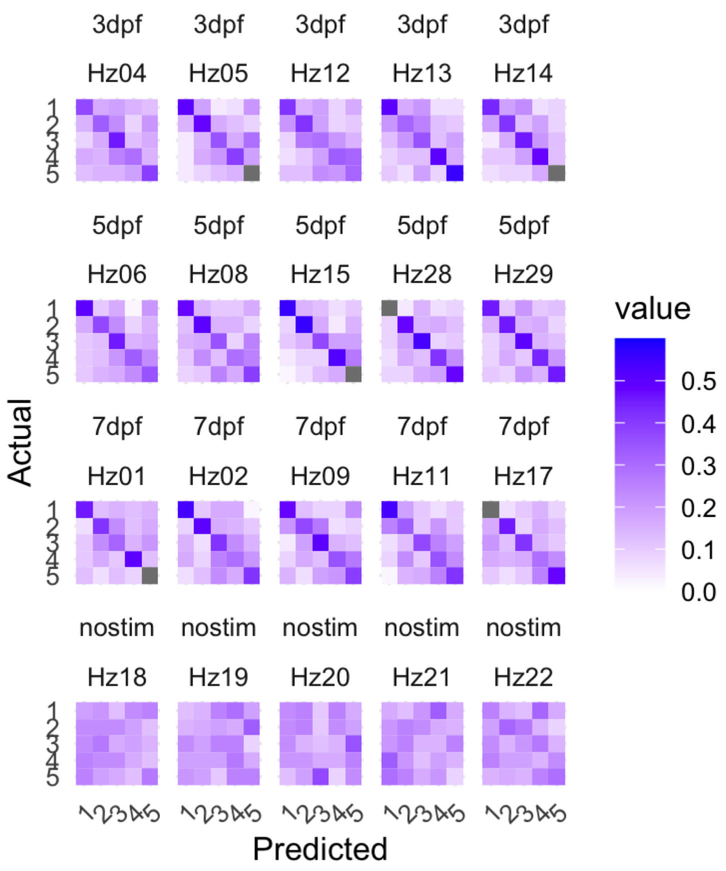


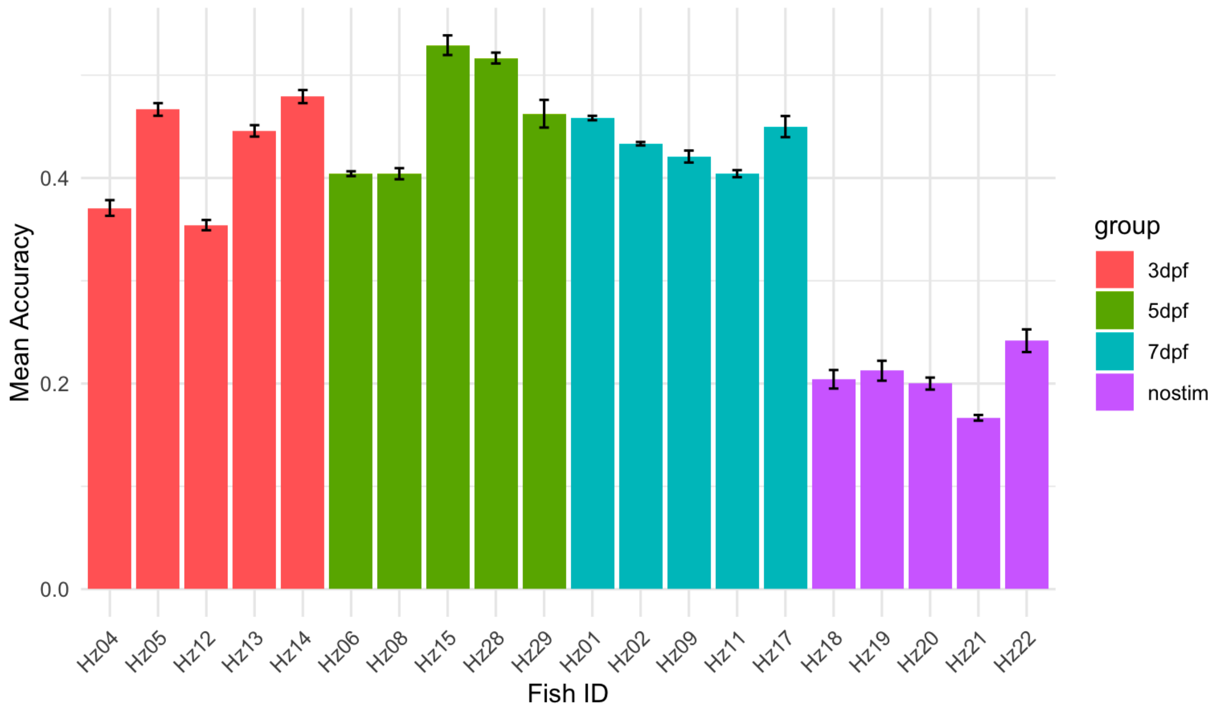

**Supplementary Table S1. Group averages of identified neurons.**

No-stimulus (negative control) group underwent normal acquisition protocol with only a visible red background and no dot stimulus.

| | Number-selective neurons | | Active neurons | |
|---|---|---|---|---|
| dpf | Average | SEM | Average | SEM |
| 3 | 766 | 306 | 13840 | 2697 |
| 5 | 796 | 101 | 16747 | 1254 |
| 7 | 550 | 108 | 16650 | 1989 |
| 7 (No-Stimulus) | 6 | 1 | 13689 | 634 |

**Supplementary Table S2. Total amount of neurons identified in the whole brain.**

Entries show 5 samples for each condition.

| Sample id | Condition | All neurons | 1-tuned | 2-tuned | 3-tuned | 4-tuned | 5-tuned |
|---|---|---|---|---|---|---|---|
| hz04 | 3dpf | 18104 | 1043 | 54 | 2 | 2 | 6 |
| hz05 | 3dpf | 14981 | 2181 | 51 | 9 | 6 | 4 |
| hz12 | 3dpf | 2346 | 362 | 1 | 0 | 0 | 0 |
| hz13 | 3dpf | 14467 | 834 | 10 | 2 | 2 | 5 |
| hz14 | 3dpf | 19303 | 1731 | 135 | 5 | 0 | 2 |
| hz06 | 5dpf | 20603 | 925 | 120 | 5 | 3 | 5 |
| hz08 | 5dpf | 19598 | 478 | 40 | 0 | 0 | 2 |
| hz15 | 5dpf | 15221 | 877 | 74 | 102 | 4 | 1 |
| hz28 | 5dpf | 13613 | 539 | 19 | 27 | 6 | 5 |
| hz29 | 5dpf | 14701 | 669 | 21 | 50 | 5 | 3 |
| hz01 | 7dpf | 23980 | 599 | 8 | 57 | 4 | 6 |
| hz02 | 7dpf | 16359 | 436 | 15 | 16 | 9 | 8 |
| hz09 | 7dpf | 14874 | 564 | 18 | 14 | 1 | 2 |
| hz11 | 7dpf | 17769 | 775 | 63 | 20 | 1 | 3 |
| hz17 | 7dpf | 10269 | 90 | 26 | 11 | 2 | 4 |
| hz18 | No-stim | 16061 | 0 | 1 | 1 | 1 | 1 |
| hz19 | No-stim | 13386 | 1 | 1 | 1 | 0 | 2 |
| hz20 | No-stim | 13247 | 1 | 1 | 2 | 3 | 2 |
| hz21 | No-stim | 11689 | 0 | 0 | 1 | 2 | 0 |
| hz22 | No-stim | 14062 | 1 | 0 | 2 | 0 | 7 |

**Supplementary Table S3. Total amount of neurons identified in the forebrain.**

Entries show 5 samples for each condition.

| Sample id | Condition | Forebrain | 1-tuned | 2-tuned | 3-tuned | 4-tuned | 5-tuned |
|---|---|---|---|---|---|---|---|
| hz04 | 3dpf | 3468 | 303 | 7 | 2 | 2 | 0 |
| hz05 | 3dpf | 2363 | 776 | 5 | 2 | 2 | 2 |
| hz12 | 3dpf | 220 | 58 | 1 | 0 | 0 | 0 |
| hz13 | 3dpf | 3671 | 179 | 2 | 0 | 2 | 3 |
| hz14 | 3dpf | 3643 | 714 | 32 | 1 | 0 | 1 |
| hz06 | 5dpf | 7095 | 596 | 4 | 0 | 1 | 1 |
| hz08 | 5dpf | 6546 | 225 | 0 | 0 | 0 | 2 |
| hz15 | 5dpf | 3495 | 355 | 14 | 2 | 0 | 0 |
| hz28 | 5dpf | 4925 | 304 | 6 | 5 | 2 | 4 |
| hz29 | 5dpf | 4871 | 257 | 3 | 0 | 1 | 3 |
| hz01 | 7dpf | 8046 | 363 | 4 | 2 | 2 | 3 |
| hz02 | 7dpf | 4160 | 82 | 3 | 0 | 3 | 4 |
| hz09 | 7dpf | 6296 | 266 | 1 | 1 | 1 | 0 |
| hz11 | 7dpf | 5292 | 293 | 2 | 2 | 0 | 0 |
| hz17 | 7dpf | 3788 | 54 | 0 | 0 | 0 | 3 |
| hz18 | no_stim | 6949 | 0 | 1 | 1 | 1 | 1 |
| hz19 | no_stim | 4493 | 1 | 1 | 1 | 0 | 2 |
| hz20 | no_stim | 4602 | 1 | 1 | 2 | 3 | 2 |
| hz21 | no_stim | 4260 | 0 | 0 | 1 | 2 | 0 |
| hz22 | no_stim | 4448 | 1 | 0 | 2 | 0 | 7 |

**Supplementary Table S4. Total amount of neurons identified in the midbrain.**

Entries show 5 samples for each condition.

| Sample id | Condition | Midbrain | 1-tuned | 2-tuned | 3-tuned | 4-tuned | 5-tuned |
|---|---|---|---|---|---|---|---|
| hz04 | 3dpf | 3665 | 660 | 29 | 0 | 0 | 0 |
| hz05 | 3dpf | 3554 | 1163 | 41 | 7 | 2 | 0 |
| hz12 | 3dpf | 741 | 288 | 0 | 0 | 0 | 0 |
| hz13 | 3dpf | 4378 | 632 | 7 | 0 | 0 | 2 |
| hz14 | 3dpf | 4392 | 927 | 94 | 1 | 0 | 1 |
| hz06 | 5dpf | 6323 | 298 | 116 | 3 | 0 | 2 |
| hz08 | 5dpf | 3502 | 203 | 40 | 0 | 0 | 0 |
| hz15 | 5dpf | 5847 | 513 | 59 | 100 | 4 | 0 |
| hz28 | 5dpf | 2603 | 171 | 12 | 20 | 2 | 0 |
| hz29 | 5dpf | 3889 | 402 | 17 | 48 | 2 | 0 |
| hz01 | 7dpf | 5936 | 166 | 2 | 52 | 2 | 3 |
| hz02 | 7dpf | 3263 | 218 | 1 | 3 | 5 | 4 |
| hz09 | 7dpf | 2468 | 242 | 13 | 8 | 0 | 2 |
| hz11 | 7dpf | 4653 | 418 | 46 | 7 | 1 | 3 |
| hz17 | 7dpf | 1614 | 29 | 21 | 11 | 0 | 1 |
| hz18 | no_stim | 1817 | 0 | 0 | 0 | 0 | 0 |
| hz19 | no_stim | 1972 | 0 | 0 | 0 | 0 | 0 |
| hz20 | no_stim | 1103 | 0 | 0 | 0 | 0 | 0 |
| hz21 | no_stim | 2620 | 0 | 0 | 0 | 0 | 0 |
| hz22 | no_stim | 1813 | 0 | 0 | 0 | 0 | 0 |

**Supplementary Table S5. Total amount of neurons identified in the hindbrain.**

Entries show 5 samples for each condition.

| Sample id | Condition | Hindbrain | 1-tuned | 2-tuned | 3-tuned | 4-tuned | 5-tuned |
|---|---|---|---|---|---|---|---|
| hz04 | 3dpf | 10971 | 80 | 18 | 0 | 0 | 6 |
| hz05 | 3dpf | 9064 | 242 | 5 | 0 | 2 | 2 |
| hz12 | 3dpf | 1385 | 16 | 0 | 0 | 0 | 0 |
| hz13 | 3dpf | 6418 | 23 | 1 | 2 | 0 | 0 |
| hz14 | 3dpf | 11268 | 90 | 9 | 3 | 0 | 0 |
| hz06 | 5dpf | 7185 | 31 | 0 | 2 | 2 | 2 |
| hz08 | 5dpf | 9550 | 50 | 0 | 0 | 0 | 0 |
| hz15 | 5dpf | 5879 | 9 | 1 | 0 | 0 | 1 |
| hz28 | 5dpf | 6085 | 64 | 1 | 2 | 2 | 1 |
| hz29 | 5dpf | 5941 | 10 | 1 | 2 | 2 | 0 |
| hz01 | 7dpf | 9998 | 70 | 2 | 3 | 0 | 0 |
| hz02 | 7dpf | 8936 | 136 | 11 | 13 | 1 | 0 |
| hz09 | 7dpf | 6110 | 56 | 4 | 5 | 0 | 0 |
| hz11 | 7dpf | 7824 | 64 | 15 | 11 | 0 | 0 |
| hz17 | 7dpf | 4867 | 7 | 5 | 0 | 2 | 0 |
| hz18 | no_stim | 7295 | 0 | 0 | 0 | 0 | 0 |
| hz19 | no_stim | 6921 | 0 | 0 | 0 | 0 | 0 |
| hz20 | no_stim | 7542 | 0 | 0 | 0 | 0 | 0 |
| hz21 | no_stim | 4809 | 0 | 0 | 0 | 0 | 0 |
| hz22 | no_stim | 7801 | 0 | 0 | 0 | 0 | 0 |

**Supplementary Table S6. Comparison of age-related change in number-selective neurons in subregions of the forebrain.**

Kruskal–Wallis test across age groups (n=5) for each region, adjusted for multiple comparisons using Bonferroni correction with alpha = 0.00625.

| Region | F value | p value |
|---|---|---|
| Dorsal Telencephalon (Pallium) | 3.42 | 0.18 |
| Eminentia Thalami | 6.74 | 0.03 |
| Hypothalamus | 8.3 | 0.02 |
| Posterior Tuberculum (Basal Part of Prethalamus and Thalamus) | 0.82 | 0.66 |
| Pretectum | 0.26 | 0.88 |
| Prethalamus (Ventral Thalamus) | 1.46 | 0.48 |
| Thalamus (Dorsal Thalamus) | 4.02 | 0.13 |
| Ventral Telencephalon (Subpallium) | 5.89 | 0.05 |

**Supplementary Table S7. Summary of confusion matrix score.**

This table presents the fraction of prediction instances using performance metrics such as precision, recall, and F1-score. Precision indicates the proportion of correctly predicted positive observations to the total predicted positives (i.e., the accuracy of positive predictions). Recall (also known as sensitivity) represents the proportion of correctly predicted positive observation to all observations in the actual class (i.e., the ability to find all relevant instances). The F1-score is the harmonic mean of precision and recall, providing a single metric that balances both concerns.

| | precision | | | | recall | | | | f1-score | | | |
|---|---|---|---|---|---|---|---|---|---|---|---|---|
| | 3dpf | 5dpf | 7dpf | No-stimulus | 3dpf | 5dpf | 7dpf | No-stimulus | 3dpf | 5dpf | 7dpf | No-stimulus |
| 1 dot | 0.53 | 0.62 | 0.55 | 0.18 | 0.44 | 0.53 | 0.53 | 0.19 | 0.47 | 0.56 | 0.53 | 0.16 |
| 2 dots | 0.41 | 0.50 | 0.44 | 0.27 | 0.40 | 0.48 | 0.39 | 0.24 | 0.40 | 0.48 | 0.41 | 0.24 |
| 3 dots | 0.36 | 0.42 | 0.39 | 0.20 | 0.38 | 0.45 | 0.41 | 0.18 | 0.36 | 0.43 | 0.41 | 0.19 |
| 4 dots | 0.44 | 0.47 | 0.37 | 0.18 | 0.41 | 0.40 | 0.37 | 0.19 | 0.42 | 0.42 | 0.37 | 0.18 |
| 5 dots | 0.44 | 0.41 | 0.43 | 0.23 | 0.49 | 0.45 | 0.46 | 0.23 | 0.46 | 0.43 | 0.45 | 0.23 |

**Supplementary Table S8. Summary of GeNEsIS parameters.**

GEnerator of Numerical ElementS Images Software (GeNEsIS) is a custom program written in Matlab to create stimuli with different numerosity and controlled or constrain physical stimuli characteristics.

| Parameter | Pixels | cm |
|---|---|---|
| Convex hull | 100 | 4.84 |
| Inter-distance | 9 | 1.98 |
| Constant radius | 1.2 | 0.05808 |
| Total area | 26 | 5.72 |
| Total perimeter | 31 | 6.82 |
| Radius variability | 0.2 | 0.044 |
| Mean inter-distance | 1.1 | 0.242 |
| Mean radius | 0.8 | 0.176 |
| Arena radius | 10 | 2.2 |
| Arena dimension (pixel) | 10 | 2.2 |
| Pixel_X screen | 1280 | 16 |
| Pixel_Y screen | 720 | 9 |

**Supplementary Table S9. Numerical values of the mean predicted accuracies and associated p-values for the SVM analysis results presented in Figure 4c and Supplementary Figure S12.**

| Type | Cohort | Fish-ID | Mean Accuracy | p-values |
|---|---|---|---|---|
| Single sample | 3 dpf | Hz04 | 0.34583333 | 0.00054776 |
| Single sample | 3 dpf | Hz05 | 0.475 | 3.3016E-05 |
| Single sample | 3 dpf | Hz12 | 0.3875 | 0.00043888 |
| Single sample | 3 dpf | Hz13 | 0.4 | 9.94E-06 |
| Single sample | 3 dpf | Hz14 | 0.49166667 | 0.00018284 |
| Single sample | 5 dpf | Hz06 | 0.4 | 3.7893E-05 |
| Single sample | 5 dpf | Hz08 | 0.3125 | 0.0255547 |
| Single sample | 5 dpf | Hz15 | 0.54166667 | 1.7645E-05 |
| Single sample | 5 dpf | Hz28 | 0.475 | 5.61E-06 |
| Single sample | 5 dpf | Hz29 | 0.46666667 | 0.00076898 |
| Single sample | 7 dpf | Hz01 | 0.42083333 | 4.9836E-06 |
| Single sample | 7 dpf | Hz02 | 0.45416667 | 4.872E-05 |
| Single sample | 7 dpf | Hz09 | 0.375 | 0.00169384 |
| Single sample | 7 dpf | Hz11 | 0.42083333 | 0.00012002 |
| Single sample | 7 dpf | Hz17 | 0.375 | 0.01435455 |
| Single sample | No-stimulus | Hz18 | 0.20416667 | 0.91080665 |
| Single sample | No-stimulus | Hz19 | 0.2125 | 0.74692699 |
| Single sample | No-stimulus | Hz20 | 0.2 | 1 |
| Single sample | No-stimulus | Hz21 | 0.16666667 | 0.13817995 |
| Single sample | No-stimulus | Hz22 | 0.24166667 | 0.32899331 |
| Mean of cohort | 3 dpf | n/a | 0.42 | 0.00132274 |
| Mean of cohort | 5 dpf | n/a | 0.43916667 | 0.00351968 |
| Mean of cohort | 7 dpf | n/a | 0.40916667 | 0.00016236 |
| Mean of cohort | No-stimulus | n/a | 0.205 | 0.69908009 |

**Supplementary Table S10. Key resources table.**

| Reagent type (species) or resource | Designation | Source or reference | Identifiers |
|---|---|---|---|
| Genetic reagent (*Danio rerio*) | Zebrafish: Tg(elavl3:H2B::jGCa MP7f) | (Dana et al., 2019; Yang et al., 2022), gift from David Prober | RRID:Addgene_1044 88 |
| Python Library | Analysis tools: ANTs | (Avants et al., 2009) | https://github.com/AN TsX/ANTs |
| Python library | Analysis tools: NuMan | This work | https://github.com/Le monJust/numan |
| Python library | Analysis tools: numan_plus | This work | https://github.com/Mir koZanon/numan_plus |
| Python library | Analysis tool: seaborn | (Waskom, 2021) | https://seaborn.pydat a.org/index.html |
| Python library | Data management: VoDEx | (Nadtochiy et al., 2023) | https://github.com/Le monJust/vodex |
| Python library | Cell Segmentation | (Giovannucci et al., 2019) | https://github.com/flat ironinstitute/CaImAn |
| Python library | Stimuli presentation: PsychoPy | (Peirce et al., 2019) | https://psychopy.org/i ndex.html |
| Software/Python Library | Image analysis toolkit: ITK-SNAP | (Yushkevich et al., 2006) | http://www.itksnap.or g/ |
| Software | Microscope GUI, μManager | (Edelstein et al., 2010a) | https://micro-manager.org |
| Software | Stimuli generation: GeNEsIS | (Zanon et al., 2022) | https://github.com/Mir koZanon/GeNEsIS |
| Figshare dataset | Dataset: json files of segmented neurons | This work | https://doi.org/10.608 4/m9.figshare.27676 197.v3 |

**Supplemental Results: Peristimulus response of all identified number-selective neurons.**

The results are contained in a zipped folder at the following online location:
https://figshare.com/articles/dataset/Peristimulus_event_of_single_neuron_activation/30174013?file=58121995

The folder contains 20 pdf files, each one corresponds to a single experimental fish. There are 5 samples for each of the age cohorts (3, 5, and 7 dpf), and 5 samples for the 'No-stimulus' control at 7 dpf. File names have strings "3dpf", "5dpf", or "7dpf" for identification. The beginning of each file name has the experimental date identifier (e.g. "20230703"), followed by the fish sample identifier (e.g. "Hz12"). Fish Hz18, Hz19, Hz20, Hz21, Hz22 are the 'No-stimulus' control fish, for which the same experimental pipeline was applied but without dots presentation.

In each pdf file, the peristimulus response of all identified number-selective neurons, for that particular fish sample, are shown. Each number neuron is represented by an image panel that spans the width of the page, with a text string at the top of the image panel that provides identifying information, including the string "num_1" or "num_4" that designates that the cell is identified as a 1-tuned or 4-tuned neuron, for example. Cells are arranged according to their CaImAn-derived number identifier. To identify cells that are tuned to a specific numerosity, readers could do a document search for "num_1", "num_2", and so on.

The image panel for each cell (an example of which is shown below) contains the information similar to the peristimulus responses shown in Fig. 2b of the Main Text, but with the difference that the responses are shown with SEM error bars, averaged over all experimental trials. In each panel, as shown below, the gray bars mark the time window of the presented numerosity stimuli, which varied from 1 to 5, going from left to right. Tick marks on the horizontal axis represent seconds. The vertical axis represents ΔF/F. In the example below, the number neuron is cell #2042, identified as a 3-tuned cell, located at XYZ location as indicated (in the reference frame of the template brain that all data from the same age cohort are registered to).

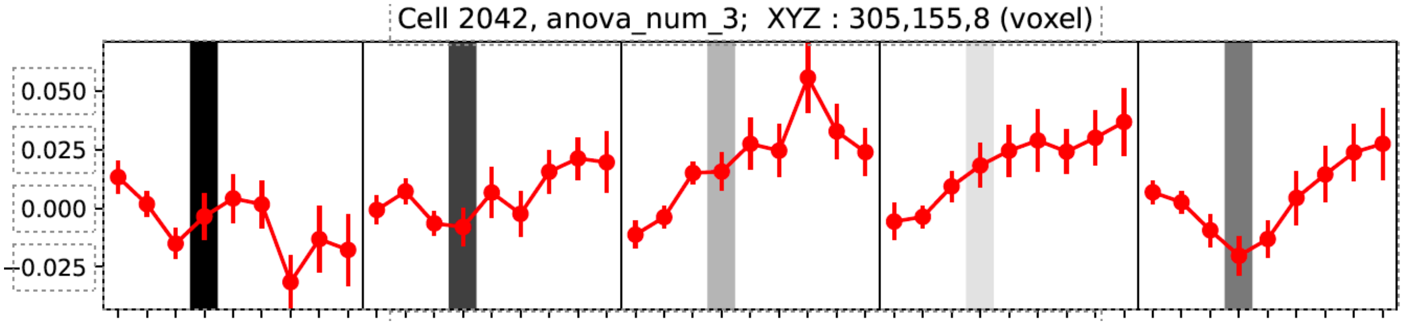
